# Biogeography and the evolution of acoustic communication in the polyploid North American gray treefrog complex

**DOI:** 10.1101/2023.04.03.535469

**Authors:** William W. Booker, Emily Moriarty Lemmon, Alan R. Lemmon, Margaret B. Ptacek, Alyssa T. B. Hassinger, Johannes Schul, H. Carl Gerhardt

**Affiliations:** Department of Biological Science, Florida State University, Tallahassee, FL 32306-4295; Department of Genetics, University of North Carolina, Chapel Hill, NC 27514; Department of Scientific Computing, Florida State University, Tallahassee, FL 32306-4192; Department of Biological Sciences, Clemson University, Clemson, SC 29634; Department of Evolution, Ecology, and Organismal Biology, Ohio State University, Columbus, OH 43210; Division of Biological Sciences, University of Missouri, Columbia, MO 65211-7400

**Keywords:** Biogeography, Polyploid, Acoustic Communication, Evolution, Anurans, Comparative Methods

## Abstract

After polyploid species are formed, interactions between diploid and polyploid lineages may generate additional diversity in novel cytotypes and phenotypes. In anurans, mate choice by acoustic communication is the primary method by which individuals identify their own species and assess suitable mates. As such, the evolution of acoustic signals is an important mechanism for contributing to reproductive isolation and diversification in this group. Here, we estimate the biogeographic history of the North American gray treefrog complex, consisting of the diploid *Hyla chrysoscelis* and the tetraploid *Hyla versicolor*, focusing specifically on the geographic origin of whole genome duplication and the expansion of lineages out of glacial refugia. We then test for lineage-specific differences in mating signals by applying comparative methods to a large acoustic data set collected over 52 years that includes *>*1500 individual frogs. Along with describing the overall biogeographic history and call diversity, we found evidence that the geographic origin of *H. versicolor* and the formation of the midwestern polyploid lineage are both associated with glacial limits, and that the southwestern polyploid lineage is associated with a shift in acoustic phenotype relative to the diploid lineage with which they share a mitochondrial lineage. In *H. chrysoscelis*, we see that acoustic signals are largely split by Eastern and Western lineages, but that northward expansion along either side of the Appalachian Mountains is associated with further acoustic diversification. Overall, results of this study provide substantial clarity on the evolution of gray treefrogs as it relates to their biogeography and acoustic communication.

## Introduction

Identifying the mechanisms that determine the spatial organization of species and their subpopulations is critical to understanding the speciation process. Although generally described as a fixed entity, a species’ range is in constant flux due to a complex interaction of abiotic and biotic forces (MacArthur 1972). Exactly how these forces interact to affect this range is, however, unclear. Kirkpatrick and Barton (1997) first formalized our understanding of range dynamics by demonstrating that a species’ range can be limited due to differential selective pressures across an environmental gradient and gene flow along that gradient. In the time since this seminal work, numerous theoretical and empirical studies have shown just how complex the evolution of a species’ range can be and the myriad forces that are at play (see Angert et al. 2020).

For polyploid taxa, understanding the dynamics that shape a species range can provide insights into how polyploids are formed and persist. In hybrid allopolyploids, where ploidy is elevated through the combination of genomes from two or more species, the dynamics regulating the proximity and interactions of related species are directly related to the likelihood that an allopolyploid species can form. Moreover, the processes that shape the number and identity of close relatives will determine the composition of polyploid complexes, which often have multiple polyploid species with varying ploidies and mixtures of subgenomes (Otto and Whitton 2000; Gregory and Mable 2005). Range dynamics play another critical part in polyploid species formation, particularly for autopolyploid species formed via whole genome duplication (WGD) of a single species, because newly formed polyploids may require unoccupied habitat to become established if they are too ecologically similar to their progenitors (Levin 1975; Parisod et al. 2010). Conversely, some evidence suggests polyploids may have a unique advantage in expanding into recently deglaciated and novel habitats (Comai 2005; Brochmann et al. 2004; Van de Peer et al. 2017; Novikova et al. 2020, 2018). The precise reason for this advantage is not entirely clear, but it may be related to the adaptive potential to novel environments due to the greater effective genic mutation rate (Otto 2007) and relaxed constraint on genes as a result of their functional redundancy (Lynch and Conery 2000; Lynch and Force 2000; Gout and Lynch 2015; Adams and Wendel 2005), the increased complexity of their gene regulatory networks enabling more diverse reconfigurations (Hegarty and Hiscock 2007; Freeling and Thomas 2006; Fusco et al. 2010; Yao et al. 2019), glacial cycles driving secondary contact (Stebbins 1984) and fixing high levels of heterozygosity in allopolyploids (Stebbins 1985; Brochmann et al. 2004), or in many cases because newly formed polyploids often switch to asexual reproduction (Comai 2005; Miller and Venable 2000) and may expand faster by ensuring reproductive success in low density populations along the expansion front (Baker 1967; Zeitler et al. 2022; Pannell and Barrett 1998). In this context, the fluctuation of glacial cycles may have been particularly important for the establishment of polyploids both by continually driving potential parental species into refugia facilitating their formation and providing advantageous conditions in which a polyploid species can persist.

Exactly how range dynamics and species interactions influence the specific mechanisms of polyploid evolution are, however, poorly understood. Although postzygotic isolation is high between species with different ploidy, there is little empirical evidence supporting the prevailing idea that “instant speciation” occurs when polyploids are formed (Barker et al. 2016). Additional doubt is raised by the fact that introgression after polyploidization is common (Marhold and Lihová 2006; Bogart and Bi 2013). Werth and Windham (1991) introduced the compelling idea that genomic restructuring of polyploid species (e.g. differential silencing and gene loss on duplicated homeologs) can result in incompatibilities between divergent populations and act as a mechanism for speciation. Support for this hypothesis in nature, however, is scarce (Muir and Hahn 2015; Li et al. 2021)—potentially due to the sensitivity of the outcome of the process to particular properties of the taxonomic groups that have been studied (Li et al. 2021).

The phenotypic and ecological consequences of polyploidization are also not well understood. Contributing the most to our understanding are studies of synthetic polyploids, which show that changes occurring immediately after polyploidization often have significant effects on morphology, reproduction, physiology, and ecological fitness (Bretagnolle and Lumaret 1995; Keller and Gerhardt 2001; Husband et al. 2008, 2016; Maherali et al. 2009; Baldwin and Husband 2011; Ramsey 2011; Oswald and Nuismer 2011; Tucker and Gerhardt 2012; Griffin et al. 2012; Martin and Husband 2012; Porturas and Segraves 2020). Research in gray treefrogs has shown that reproductive isolation mechanisms can immediately form upon polyploid formation, as a result of cell size increases that lead to corresponding changes in acoustic signals (Keller and Gerhardt 2001) and female selectivity based on these call changes (Tucker and Gerhardt 2012). These changes do not require any differences in genomic sequence, but rather result from increases in cell size and decreases in cell numbers in polyploids that include tissues involved in producing calls and auditory processing (Gerhardt et al. 2022). As such, evidence from these studies suggest that mechanisms of sexual selection, here female choice of male acoustic signals, can provide a conduit with which polyploids are rapidly differentiated from their progenitors facilitating their establishment and subsequent expansion.

The North American gray treefrog complex, composed of the diploid *Hyla chrysoscelis* and the tetraploid *Hyla versicolor*, provides a particularly intriguing opportunity to understand the role that range dynamics and species interactions play in shaping diversity during polyploid speciation. These two species are widespread across eastern North America, with a range that encompasses all land east of southern Texas and north to southern Canada, and the two species have considerable overlap in sympatry as well as allopatric areas and disjunct populations. Due to this mosaic distribution and its reticulate nature, the origins of this complex have been debated for more than four decades (Maxson et al. 1977; Ptacek et al. 1994; Holloway et al. 2006; Bogart and Bi 2013; Bogart et al. 2020; Ralin 1978). The most recent and comprehensive study of these origins indicate that the tetraploid originated through a single autopolyploid whole genome duplication event, but that several polyploid lineages subsequently resulted from repeated reticulations with divergent diploid populations and mitochondrial capture (Booker et al. 2022). Furthermore, no study to date has addressed how the large diversity of acoustic signals across lineages (e.g., Gerhardt 1999, 2005, 2013; Holloway et al. 2006) has originated in the context of biogeography.

Here, we aim to unravel the biogeography and the patterns of acoustic evolution in the North American gray treefrog complex. Using phylogenies estimated from hundreds of nuclear loci and whole mitochondrial genomes from a previous study (Booker et al. 2022), we build upon this work describing the origins and systematics of this group by estimating the biogeographic history of polyploid formation. In estimating this history, we specifically test the hypothesis that the origin of WGD and other lineage formations are associated with the limits of the Laurentide ice sheet during Quaternary periods of glacial oscillation. We then combine these analyses with acoustic data from 1,613 frogs gathered over 52 years of field sampling to conduct a species complex-level analysis of acoustic evolution using phylogenetic comparative methods and normal mixture models to test for lineage specific evolution of acoustic signals within both diploids and tetraploids. Finally, we outline a comprehensive history of this complex by interpreting our observations of acoustic evolution in light of our estimates of biogeography.

## Methods

### Phylogenetic Data

The phylogenetic data used for this study were obtained from publicly available data generated for a previous study by Booker et al. (2022). Briefly, the phylogenies used were generated from sequence data collected using Anchored Hybrid Enrichment (Lemmon et al. 2012). Nuclear phylogenies for *H. chrysoscelis* were estimated under maximum likelihood (Stamatakis 2014) using a concatenated alignment of 244 loci (374,891 total sites) and 82 individuals (including *H. avivoca*, *H. arenicolor*, *H. andersonii*, and *H. femoralis* as outgroup taxa). Whole mitochondrial phylogenies were generated by first mapping bycatch reads from sequencing runs originally targeting nuclear sequences to previously generated mitochondrial genomes of a close relative, aligning the newly assembled mitochondrial genomes, and then using BEAST (Drummond and Rambaut 2007; Bouckaert et al. 2014) to estimate the phylogenetic relationships. Coalescent timing was conducted alongside phylogenetic estimation of mitochondrial genomes using previously estimated molecular clocks of a relative species. The final sequence length of mitochondrial alignments was 15,834 sites and the original phylogeny was generated using 117 individuals (including *H. avivoca*, *H. arenicolor*, *H. andersonii*, and *H. femoralis* as outgroup taxa; See Booker et al. 2022 for full details).

### Biogeographic Analyses

To estimate biogeographical patterns and test hypotheses under a likelihood based framework, we used the program PhyloMapper from Lemmon and Lemmon (2008). Briefly, PhyloMapper works by using a phylogenetic tree and latitude and longitude coordinates of the tips to model the geographic location of the tree nodes and dispersal of lineages from those nodes. The analyses we conducted broadly aimed to: (1) identify the origin of *H. versicolor* lineages and the estimated location of WGD, (2) identify potential glacial refugia in *H. chrysoscelis*, and (3) estimate the direction and routes of expansion for both species. Because we were interested in making inferences for individual species on their own, we generated new trees for the two species individually for our analyses with the same alignment and parameters used to generate the full tree from Booker et al. (2022). For the individual species trees, we estimated phylogenetic relationships using the full alignment, only removing individuals of non-focal species lineages that did not form a monophyletic group inside a focal species lineage (i.e. removing *H. versicolor* individuals in the SW/CSW mitochondrial clade for *H. chrysoscelis* analyses; light blue, Fig. 1; Supp. Fig. 1), as the multi-species relationships of this group are likely due to phylogenetic estimation error as a result of the low level of sequence diversity within the group. All other non-focal species individuals were then pruned from the analyzed tree. We used the mitochondrial tree because it matches the PhyloMapper’s assumption of a known genealogy better than would poorly resolved trees derived from single locus or a concatenated nuclear tree that contains a mixture of nuclear gene histories (see below where we assess the effect of phylogenetic uncertainty to test this assumption). Although using mitochondrial data alone to infer biogeography is problematic (see, Edwards 2009; Toews and Brelsford 2012), the nuclear genomic history has been investigated in depth previously (Booker et al. 2022), and as we make evident, mitochondrial introgression in this group highlights important evolutionary events that can be placed in context with the estimations from Booker et al. (2022).

**Figure 1.**
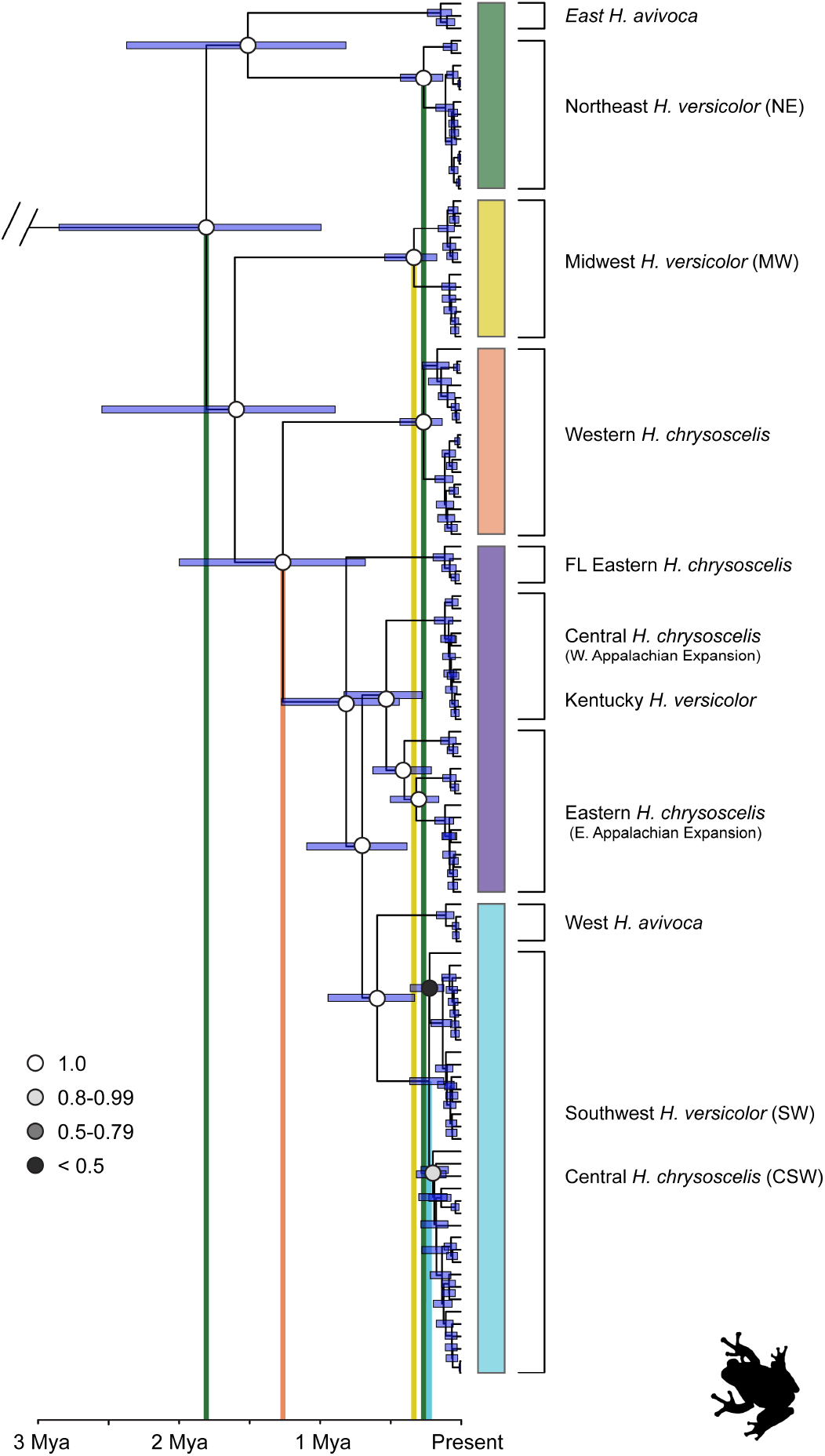
Dated whole-genome mitochondrial phylogeny estimated in Booker et al. (2022). Colored bars right of the phylogeny highlight mitochondrial clades. Circle color on nodes represent posterior values for those nodes and are only reported for branches informative for this study. From left to right, vertical bars show mean timing of coalescence for: 1) *H. avivoca*, *H. versicolor,* and *H. chrysoscelis*; 2) Eastern/Central *H. chrysoscelis* and Western *H. versicolor*, 3) all MW *H. versicolor* ; 4) all NE *H. versicolor* ; and 5) all SW *H. versicolor* and the Central (CSW) *H. chrysoscelis* with which they share a monophyletic mitochondrial clade.

To conduct our analyses we began by identifying clades of interest that have recently expanded by testing for dispersal rate (*ψ*) heterogeneity using PhyloMapper (Lemmon and Lemmon 2008), based on the mitochondrial genome phylogeny and population structure delimited in Booker et al. (2022). Dispersal rates are expected to be higher for lineages that have recently expanded (Kirkpatrick and Barton 1997; Lemmon and Lemmon 2008). Following Lemmon and Lemmon (2008), we conducted a series of nested likelihood ratio tests that compared a model containing separate dispersal rates for two or more different clades to a null model containing a single dispersal rate for the different clades. Taking a hierarchical approach, we applied this test from the shallowest clades then proceeding deeper into the tree, collapsing dispersal classes for clades that could not be distinguished significantly.

After testing for dispersal rate heterogeneity, we estimated the location of lineage ancestors and potential refugia, taking into account uncertainty due to the dispersal model and uncertainty due to mitogenome estimation. Since PhyloMapper estimates a single ancestral location by maximum likelihood, it is important to account for uncertainty due to the stochastic nature of the model (Lemmon and Lemmon 2008). To test whether the envelope of uncertainty includes the location expected under the null model (the center of the sampled points), we compared the maximum likelihood model to geographic center and alternative models using ΔAIC. To estimate the geographic center of the sampled localities, we randomized the assignment of coordinates to tips within the focal clade for 10,000 replications and averaged the maximum likelihood coordinates of the ancestor across all replications.

In addition to assessing the geographic center and maximum likelihood estimated ancestor models, we also assessed models where clades expanded out of alternate refugia when such a scenario seemed plausible. To do this, we restricted the ancestor coordinates to minimum and maximum latitudes and longitudes encompassing the proposed alternative refugium, and we estimated the maximum likelihood of each point in that grid in 0.2 coordinate increments. We then used the coordinates within that grid that had the highest maximum likelihood score for our final model test. Once all coordinates and likelihoods for the models assessed had been estimated, we assessed model probability using ΔAIC calculations where the maximum likelihood (ML) had two additional parameters compared to the the geographic center (C) or the alternative refugium (AR) models. Next, to examine the potential effect of genealogical uncertainty on the estimates, we randomly sampled 1000 trees from the posterior distribution of trees, repeated the PhyloMapper analyses, then plotted the distribution of estimated ancestral locations. Finally, we estimated migration routes and tested for non-random directionality by connecting the location of the ancestor to the location of its descendants and testing for a deviation in the distribution of connection angles from the null distribution of angles generated from our previous 10,000 randomizations.

### Acoustic Data Collection and Temperature Correction

In total, we used calls from 1182 *H. chrysoscelis* and 431 *H. versicolor* individuals (12,457 total calls, mean 7.72 calls/individual) recorded between 1963 and 2015. Recordings of advertisement calls were made with directional microphones (Sennheiser [Wedermark, Germany] models 415, 815, ME80) and various tape recorders (Nagra [Cheseaux, Switzerland] models IV-SR, IV-S; Stellavox [Geneva, Switzerland] reel-to-reel; Sony [Tokyo, Japan] Walkman Professional cassette, Walkman DAT; Marantz [Cumberland, RI, USA] PMD671 digital). The specifications of all recording equipment were more than adequate to resolve the values of the fine-scale temporal properties of the calls to within 1% or less. For each recording, the microphone was placed about 25–50 cm from the calling male and oriented directly toward it.

Because pulse rate increases linearly with the temperature of the calling frog (r *>*0.95; Gerhardt 1978, 2005), we recorded the most accurate temperature possible given the situation. A quick-reading thermometer (Schultheis, Queens, New York) was used to obtain a cloacal or surface temperature of most recorded frogs. Most recordings were followed by an attempt–usually successful–to obtain a cloacal temperature. If there was a strong chance that handling or escape and recapture changed the frog’s temperature, temperature readings of the calling site (air or water) were made. Temperature measurements are always subject to error regardless of the method and equipment because humidity and frog calling site vary such that wet-bulb readings will be more reliable in some situations and dry-bulb in others. As a final check, we compared the temperature estimates by site/night to identify and correct possible errors.

The spectral properties of the calls were analyzed using a Kay (Pine Brook, NJ) 210 5500 DSP Sona-Graph. This instrument also digitized the signals (12-bit, 10 kHz sampling rate) for rapid and efficient analysis of the temporal properties using software written by G. Klump, D. Polete and J. Brown. The software computed the mean pulse rate (PR, pulses/second) of each call and the grand mean of pulse rate and number of pulses per call (PN, pulse number) for that individual across all calls (*>* 2 calls). We used two calls as a minimum, because previous research has shown the values of pulse rate for just two calls are informative because of the extreme stereotypy of this property within males (Gerhardt 1991). The grand means of pulse rate within each population with a reasonable sample size were regressed against effective (usually body; see above) temperature after outliers had been eliminated by criteria discussed in Leroy and Rousseeuw (1987)). These coefficients, in turn, were used to estimate the mean pulse-repetition rate of each population at 20°C, which is near the middle of the temperature range (about 16–25°C) at which most breeding takes place. Temperature-correction of pulse rate for populations for which there was a small sample size was based on nearby populations of the same lineage for which there was an adequate sample size. The first 1-3 pulses of each call were not reported because the low amplitude of these pulses made their identification unreliable. We chose to use pulse rate for this analysis, because this character is important for species discrimination in gray treefrogs (Gerhardt 2005), shows evidence of reproductive character displacement (Gerhardt 1994, 2013), and is directly affected by polyploidy (Keller and Gerhardt 2001; Tucker and Gerhardt 2012). Similarly, we chose to use pulse number because this character interacts with pulse rate to affect call duration (PN/PR), and call duration influences intraspecific female preferences and is a reliable indicator of heritable genetic quality (Welch et al. 1998; Welch 2003). Finally, in comparison to other systems, pulse rate and pulse number appear to be important in sexual selection in anurans with similarly structured calls generally (Lemmon 2009; Lemmon and Lemmon 2010). Note that the number of pulses produced by a male, though not affected by temperature, is somewhat plastic since males can modify the pulse number in response to background noise, chorus density or the calls of nearby individuals (Wells and Taigen 1986; Gerhardt et al. 1996; Love and Bee 2010).

### Acoustic Communication Analyses

We used the R package PhylogeneticEM version 1.4.0 to test for convergence of call characters and to determine the pattern of selective regimes for the acoustic data in a phylogenetic context (Bastide et al. 2018a). Briefly, PhylogeneticEM works by estimating the likelihood of trait evolution along a tree under various models, taking an ultrametric tree and trait values of the tree tips as its input. To determine the number of selective regimes, the software models trait evolution as an Ornstein-Uhlenbeck (OU) process where stabilizing selection pulls trait values towards some optimum, and shifts are modeled along the tree as clade-specific optima. Users specify the range of possible optima, and PhylogeneticEM estimates the maximum likelihood solution for each optimum value. The number of optima is then determined by a penalized likelihood criterion to avoid overfitting. Additionally, to avoid issues of independence between traits, PhylogeneticEM allows the use of a scalar OU model of trait evolution, and this model was used for the present analysis.

We investigated the pattern of acoustic evolution with respect to pulse rate (temperature corrected to 20°C) and pulse number, two traits that are critical for species and mate recognition in gray treefrogs and other frog species (Gerhardt and Huber 2002). To conduct comparative analyses, we used a total of 538 individuals across 45 populations, and allowed for PhylogeneticEM to estimate values for populations lacking acoustic data. Population data was calculated as the average across all individuals within the same county or an adjacent county if acoustic data were not available for that tip location. In one case, Phelps Co. Missouri, we had acoustic and sequence data matched in two separate populations, and both were included in this analysis. We used the concatenated nuclear tree of *H. chrysoscelis* generated from Booker et al. (2022) to conduct this analysis because nuclear species trees reflect the true speciation history more accurately than mitochondrial trees, and therefore better reflect the evolutionary history of calls. However, these results are also considered in context with our biogeographic analyses to illustrate the relationship between biogeographic history and call evolution in gray treefrogs.

One issue with using comparative methods such as PhylogeneticEM is that they generally assume there is no gene flow across the phylogeny. Given previous results (Booker et al. 2022), we know this assumption is violated. Although there are comparative methods that incorporate gene flow (Bastide et al. 2018b), these methods are intractable for large trees such as the one used here. To attempt to verify our comparative analyses, we also modeled calls as a mixture of normal distributions using normal mixture models (NMM; Peel and MacLahlan 2000). These models attempt to model the distribution of data not as a single distribution but instead as a mixture of one or more normal distributions. NMMs have been previously used to test hypotheses of species delimitation with phenotypic data (Cadena et al. 2018), and should therefore provide a more conservative alternative estimate of the number of selective regimes proposed by PhylogeneticEM. We used the R package mclust version 5.4.9 (Scrucca et al. 2016) to model the number of components (i.e. distributions) that best fit the distribution of calls represented by pulse rate and pulse number (log transformed). We estimated the number of components between 1 and 10 as the model and number of components that maximized the Bayesian Information Criterion (BIC), assigning each individual to a specific component. To more directly test the estimation of regimes by PhylogeneticEM, we fit NMMs using the populations from the PhylogeneticEM analysis only, attempting to fit models with three components delimited by the regimes determined by PhylogeneticEM *H. chrysoscelis*, or two components where two regimes are collapsed into each other and the third remains distinct (e.g. E + C and W as the two components).

Phylogenetic comparative analyses were only conducted for *H. chrysoscelis*, because a similar analysis for the polyploid *H. versicolor* would heavily violate the assumptions of these analyses. Although no estimated phylogeny represents the exact true evolutionary history of its taxa, the *H. chrysoscelis* concatenated nuclear phylogeny is largely representative of the evolutionary history of that group, as is shown from multiple lines of evidence by Booker et al. (2022). For *H. versicolor*, however, there are several issues with using any of the estimated phylogenies. First, neither the concatenated nuclear nor mitochondrial phylogenies accurately capture the highly reticulate nature of *H. versicolor* in this complex on their own, and accurate inference of this history requires considering both these phylogenies alongside inferences into the processes that generated their topologies (e.g. migration and speciation model tests, Booker et al. 2022). Second, evidence suggests that this introgression may have had a large effect on the acoustic signals for this lineage (see Acoustic Analysis of *H. versicolor* Results). Although recent methods allow comparative analyses to be conducted in a phylogenetic network (Bastide et al. 2018b), these methods are not tractable for large data sets.

Because we lacked the ability to explicitly test models of call evolution in a phylogenetic context for *H. versicolor*, we instead opted to test for differences in calls between mitochondrial lineages themselves. These lineages represent the original WGD population (Northeast: NE) and two populations which exhibit introgressed mitochondria from extinct and extant populations of *H. chrysoscelis* (Southwest: SW; Midwest: MW) (Fig. 1) (Booker et al. 2022). We used mitochondrial lineages, because these mitochondrial lineages delineate major evolutionary events in *H. versicolor*, and previous work along with a preliminary view of the data show (Fig. 3a) a strong overlap between the geographic distribution of the SW mitochondrial lineage and the distribution of calls with a faster pulse rate (Holloway et al. 2006). To conduct this analysis, we used call measurements for 432 individuals from 61 populations across the range of *H. versicolor* to test for differences between phylogeographic groups. Similar to our analysis in *H. chrysoscelis*, we chose to test for differences in pulse rate (corrected to 20°C) and pulse number, using the average value for each sampled population in our analyses.

For each test between *H. versicolor* lineages, we generated a test statistic that was the mean character value of one lineage divided by the mean of the second lineage. We then conducted randomization tests using this statistic by generating 100,000 null replicates where the lineage assignment was randomized prior to calculating the statistic. Significance was assessed as the number of null replicates that were greater than or equal to our observed statistic divided by the total number of null replicates. Because we did not have individual frog calls matched to mitochondrial genomes, we excluded individuals that were sampled near or within lineage contact zones to ensure we were testing the true differences associated with each lineage mitochondrial haplotype.

Although we were unable to use PhylogeneticEM to analyze *H. versicolor* calls, we were able to use NMMs to model the number of components similar to *H. chrysoscelis*. In addition to estimating the number of components generally, we also tested for differences between the mitochondrial lineages. For this analysis, we fit an NMM using all three lineages, excluding the contact zone, as the components of the NMM and compared this model to NMMs where two mitochondrial lineages were collapsed into one (e.g. NE + MW as one component, SW as the second).

## Results

### Biogeographic Analysis of H. versicolor

Our ML estimated location for the origin of *H. versicolor* by WGD (NE coalescence) places the polyploid’s formation near the New York and Connecticut border (Fig. 2a). This estimation places the origin of *H. versicolor* just north of the Illinoian glacial limit (Fullerton 1986) which occurred around 190–130 Ka (Curry et al. 2011) (Heavy dashed line, Fig. 2a), a period close to recent (100Ka (ABC), 262Ka (Phylogenetic) Booker et al. 2022) and past estimates (pre-Wisconsin, Blair 1965) of the original WGD event. The ML model received greater support, with moderate significance, than the geographic center model for NE *H. versicolor* from our AIC tests (ΔAIC = 1.54; Table 2), providing additional support for this origin location. This result is also robust to phylogenetic estimation error (Fig. 2a). Although dispersal tests did not detect any significant difference in rates across *H. versicolor* lineages (Table 3), *H. versicolor* had a generally higher dispersal rate (*ψ*=527.4) than most *H. chrysoscelis* lineages (*ψ*=262.4) except for those where there is evidence of more recent expansion (see Discussion)—suggesting recent expansion of *H. versicolor* across all lineages. Similarly, directionality was found to be non-random for all lineages (*p<*0.0001) providing additional support for recent expansion. The data suggest that the direction of expansion was north and south for NE *H. versicolor*, east and west in MW *H. versicolor*, and north and south in SW *H. versicolor* (Supp. Fig. 2; Supp. Fig. 3; Supp. Fig. 4).

**Figure 2.**
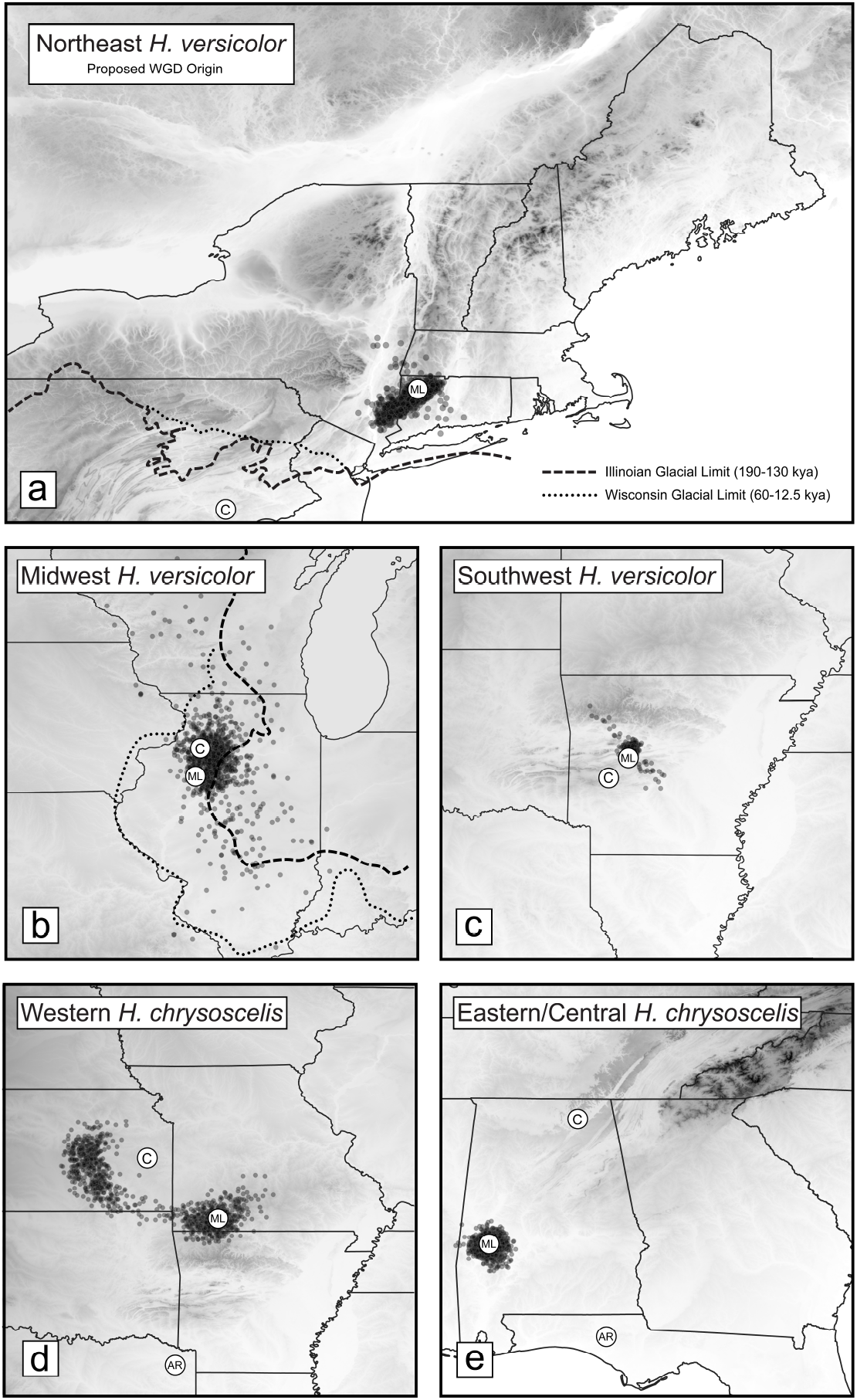
Estimated maximum likelihood (ML), geographic center (C), and Alternative Refugium (AR) ancestor locations from PhyloMapper analyses. Map shading indicates relative elevation with higher elevations darker. Opaque black circles are individual ML estimates of the ancestor location for each of the 1000 trees sampled from the posterior to assess the effect of phylogenetic uncertainty on location estimates. Heavy and light dashed lines indicate the Illinoian and Wisconsin glacial limits (respectively) from Fullerton (1986).

Although phylogenetic uncertainty resulted in somewhat reduced precision, our estimates of the geographic origins of the reticulate MW *H. versicolor* lineage placed the ancestor in northwest Illinois, in between the Illinoian and Wisconsin (approximately 60–12.5 Ka Curry et al. 2011) glacial limits (Fig. 2b; Fullerton 1986). Our ML estimate performed worse than the geographic center model (Table 2), indicating that we cannot reject the center of the range as the geographic origin. Nonetheless, the high dispersal rate and directionality suggest the population has expanded and is not at equilibrium.

Finally, though there is some dispersion due to phylogenetic uncertainty, our estimate of the origin of reticulate SW *H. versicolor* is largely within the Arkansas River Valley between the Ouachita and Boston Mountains in Arkansas (Fig. 2c). Like the MW lineage, the ML estimation model of the SW *H. versicolor* performed worse than the geographic center model (Table 2) but showed a high dispersal rate and directionality. Additionally, when we removed CSW *H. chrysoscelis* from this clade to conduct this analysis, we uncovered a significant association between geographic and genealogical proximity (*p<*0.0001) in SW *H. versicolor*. Visual inspection of the phylogeny produced after removing CSW *H. chrysoscelis* demonstrates a more plausible topology of SW *H. versicolor* with high branch support (Supp. Fig. 5), compared to the full mitochondrial tree from Booker et al. (2022), which demonstrated implausible geographic associations between SW *H. versicolor* sequences and their respective CSW *H. chrysoscelis* relatives (Fig. 1; Supp. Fig. 1). These results—in addition to the lack of contemporary introgressions observed in other *H. versicolor* outside of a single individual in an isolated Kentucky population, as well as in investigated sympatric populations (Bogart et al. 2020) and natural hybrids (Gerhardt et al. 1994; Bogart and Bi 2013)—suggest the spurious relationships seen in the full tree are due to estimation error rather than multiple introgression events.

### Biogeographic Analysis of H. chrysoscelis

Biogeographic analyses conducted using PhyloMapper suggest that the current distributions of Eastern/Central and Western nuclear genetic lineages of *H. chrysoscelis* are the result of multiple expansions from ancestral locations in southern Alabama and somewhere in the Ozarks or Great Plains around the intersection of Missouri, Arkansas, Oklahoma, and Kansas—respectively (Fig. 2d-e, Fig. 3d). For the Eastern/Central *H. chrysoscelis* lineage, our ML estimate of the ancestor location in the East Gulf Coastal Plain near the Mississippi border of southern Alabama was robust to phylogenetic estimation error, and this model outperformed the geographic center model which placed the ancestor in the Cumberland Plateau of northeast Alabama (ΔAIC = 10.70; Table 2, Fig. 2e). We found significant directionality (*p<*0.0001) and evidence for disparate dispersal rates within Eastern/Central *H. chrysoscelis* for three clades: Central *H*. *chrysoscelis* within the SW mitochondrial lineage (CSW) (*ψ*=258.09) that expanded west from Alabama, Eastern/Central *H. chrysoscelis* (*ψ*=546.47) that expanded northward on either side of the Appalachian Mountains, and a relict clade in Florida that is sister to all other Eastern/Central *H. chrysoscelis* (*ψ*=169.45) (Table 3; light blue and purple clades, Fig. 1). The phylogenetic relationship and low dispersal rate of the relict Florida clade, along with the presence of additional haplotypes from FL associated with more northern clades (Fig. 1; extra invasion seen in Supp. Fig. 6), suggest a potential stable refugium for *H. chrysoscelis* may have also existed in the Big Bend region and northern peninsula of Florida. We found little support for the hypothesis that this was the refugium for all Eastern/Central *H. chrysoscelis*, however, when we constrained the ancestor of Eastern/Central *H. chrysoscelis* to this area (Table 2). Finally, the higher dispersal rate for the broad Appalachian clade and lower dispersal rate in CSW and the relict FL *H. chrysoscelis* suggest populations in the southeastern United States have been generally more stable, with expansion north along either side of the Appalachians occurring more recently.

**Figure 3.**
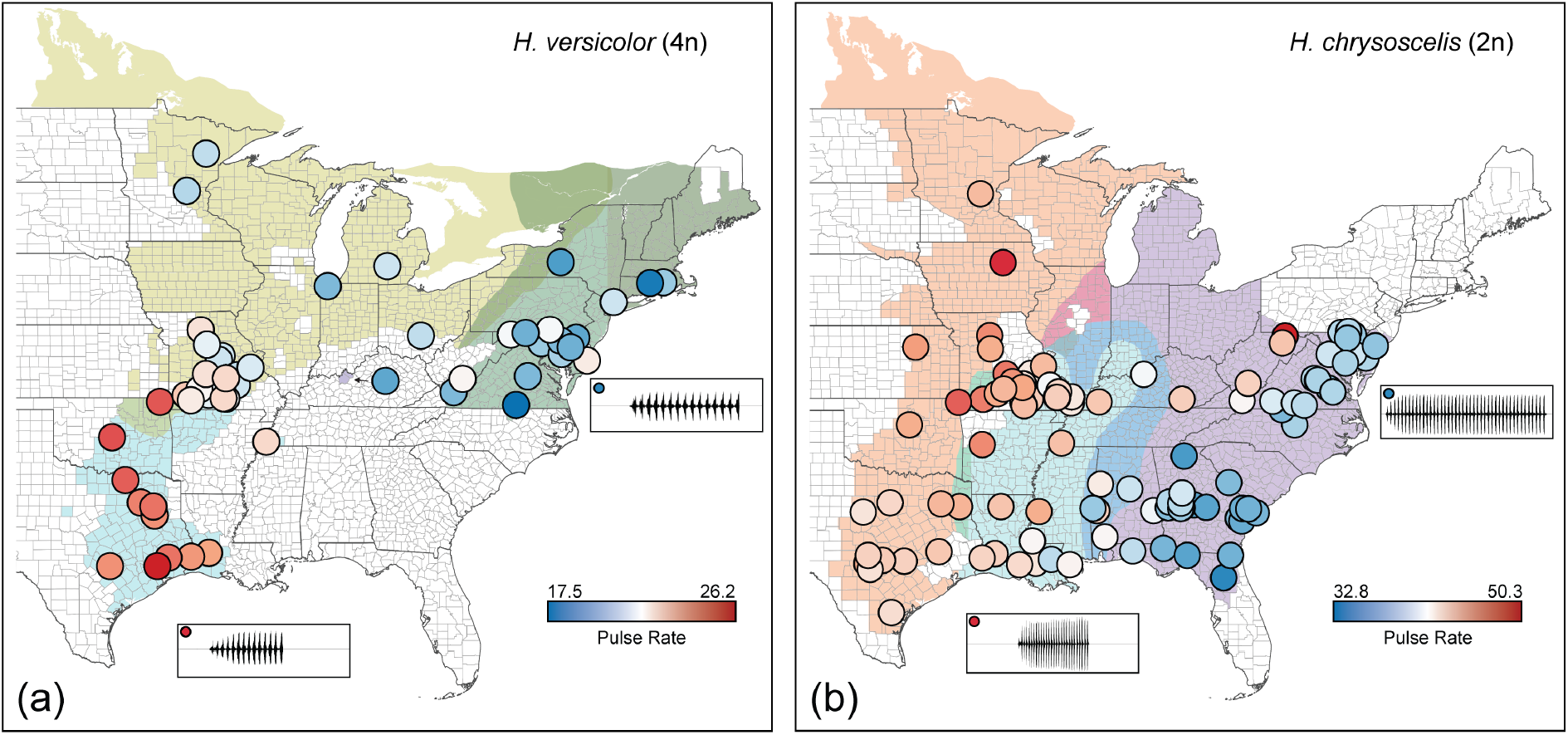
Biogeography and acoustic evolution of gray treefrogs. (a-b) Pulse rate of populations across the range of *H. versicolor* and *H. chrysoscelis*. Background colors show the estimated ranges of mitochondrial lineages from Booker et al. (2022). Circles represent a population mean colored relative to slow (blue) or fast (red) pulse rates (pulses/s). Oscillograms in inset boxes are from a single individual that represent pulse rate extremes for that species.

Our ML estimate of the Western *H. chrysoscelis* placed the lineage ancestor of this group in the Ozark Mountains of southern Missouri (Fig. 2d), but this model performed worse than the geographic center model (Table 2). We found a largely bimodal distribution of ancestor location estimates from the posterior trees when assessing for phylogenetic uncertainty (Fig. 2d), suggesting a frequent alternate topology generated when estimating this tree that would estimate a different ancestor location in eastern Kansas. The largely north-south expansion directionality (Fig. 3d, Supp. Fig. 7) was significant for this clade (*p<*0.0001), and descendants from this lineage had the highest observed dispersal rate in *H. chrysoscelis* (*ψ*=624.72) (Table 3), suggesting perhaps the most recent *H. chrysoscelis* expansion occurred in the Western lineage, a pattern consistent with the estimated early coalescent time of its extant members (Table 1). We also found some limited evidence of another refugium in eastern Texas, supported by the relationship of a *H. chrysoscelis* haplotype there with north-expanding Western *H. chrysoscelis* (Supp. Fig. 7), as well as their shared acoustic selective regime (see Acoustic Analysis of *H. chrysoscelis* Results). Similar to the FL refugium, we tested if eastern Texas was the refugium of all Western *H. chrysoscelis*, but we found very little support for this model (Table 2).

**Table 1.** Coalescent timing for clades of interest from Booker et al. (2022). Dates correspond to the phylogeny in Fig. 1.

| Clade | Fig. 1 Color | Mean | 95% CI |
| --- | --- | --- | --- |
| All *H. chrysoscelis*/*H. versicolor*/*H. avivoca* | NA | 1.80 Ma | 0.989-2.85 Ma |
| All *H. chrysoscelis* | NA | 1.26 Ma | 0.675-1.99 Ma |
| NE *H. versicolor*/East *H. avivoca* | Green | 1.51 Ma | 0.812-2.37 Ma |
| NE *H. versicolor* | Green | 0.262 Ma | 0.125-0.426 Ma |
| MW *H. versicolor* | Yellow | 0.338 Ma | 0.131-0.430 Ma |
| Western *H. chrysoscelis* | Orange | 0.262 Ma | 0.130-0.430 Ma |
| FL Eastern/Eastern/Central *H. chrysoscelis* | Purple | 0.811 Ma | 0.431-1.27 Ma |
| Eastern/Central *H. chrysoscelis* | Purple | 0.527 Ma | 0.271-0.826 Ma |
| West *H. avivoca*, SW *H. versicolor*, Central *H. chrysoscelis* (CSW) | Light Blue | 0.591 Ma | 0.326-0.939 Ma |
| SW *H. versicolor*, Central *H. chrysoscelis* (CSW) | Light Blue | 0.223 Ma | 0.012-0.360 Ma |

**Table 2.** Model likelihoods, coordinates, and probabilities for the clade ancestor location using the maximum likelihood, geographic center, or alternative refugium location estimated from PhyloMapper.

| Clade | Model lnL | Latitude | Longitude | ΔAIC |
| --- | --- | --- | --- | --- |
| **NE *H. versicolor*** |  |  |  |  |
| ML | -125.09 | 41.7533 | -73.2117 | 0.00 |
| Geo. Center | -126.86 | 39.8516 | -76.2790 | 1.54 |
| **MW *H. versicolor*** |  |  |  |  |
| ML | -124.87 | 40.7793 | -89.9611 | 1.94 |
| Geo. Center | -124.90 | 41.4129 | -89.8124 | 0.00 |
| **SW *H. versicolor*** |  |  |  |  |
| ML | -87.97 | 35.0989 | -93.2935 | 3.80 |
| Geo. Center | -88.07 | 34.4889 | -93.7771 | 0.00 |
| **E/C *H. chrysoscelis*** |  |  |  |  |
| ML | -563.35 | 32.3044 | -87.8979 | 0.00 |
| Geo. Center | -569.70 | 34.6849 | -86.2909 | 10.70 |
| Florida AR | -565.58 | 30.6000 | -85.8000 | 2.46 |
| **W *H. chrysoscelis*** |  |  |  |  |
| ML | -202.81 | 36.6807 | -93.3178 | 1.76 |
| Geo. Center | -202.93 | 38.1889 | -94.7935 | 0.00 |
| Texas AR | -222.69 | 29.0000 | -94.8000 | 37.76 |

**Table 3.** Dispersal rates (*ψ*), change in log likelihood, and *p*-values from chi-squared tests (df=1) of dispersal classes for each clade. ΔlnL and *p*-values are from tests of a unique dispersal class for the given clade against a model where that clade is collapsed into the less-indented clade above it (e.g. W *H. chrysoscelis* into All *H. chrysoscelis*). Bold clades indicate clades with unique dispersal classes.

| Clade | $\psi$ | $\Delta\ln L$ | $p$-value |
| --- | --- | --- | --- |
| All *H. versicolor* | 527.42 | — | — |
| NE *H. versicolor* | 494.02 | 0.79 | 0.374 |
| W *H. versicolor* | 543.77 | 0.10 | 0.756 |
| MW *H. versicolor* | 588.14 | 0.94 | 0.332 |
| SW *H. versicolor* | 484.06 | 0.13 | 0.714 |
| All *H. chrysoscelis* | 262.24 | — | — |
| **E/C *H. chrysoscelis*** | 348.50 | — | — |
| Relict FL ***H. chrysoscelis*** | 169.45 | 9.78 | 0.0018 |
| Appalachian/CSW *H. chrysoscelis* | 333.98 | 2.07 | 0.1505 |
| **Appalachian *H. chrysoscelis*** | 546.47 | 19.91 | <0.0001 |
| E Appalachian *H. chrysoscelis* | 544.18 | 0.72 | 0.3955 |
| W Appalachian *H. chrysoscelis* | 554.80 | 0.72 | 0.3956 |
| **CSW *H. chrysoscelis*** | 258.09 | 18.50 | <0.0001 |
| **W *H. chrysoscelis*** | 624.72 | — | — |
| Northern W *H. chrysoscelis* | 660.97 | 0.11 | 0.7398 |
| Southern W *H. chrysoscelis* | 621.28 | 0.00 | 0.9606 |

### Acoustic Analysis of H. versicolor

We found significant variation in pulse rate of acoustic signals among the *H. versicolor* lineages (*p<*0.001; SW = 22.73, MW = 20.57, NE = 19.22; 3a). The greatest difference in pulse rate was observed between SW and NE *H. versicolor* with a pulse rate difference of 3.51 pulses/second (95%CI = 2.79–4.23). The least differentiated lineages were MW and NE lineages with a difference of 1.35 pulses/second (95%CI = 0.54–2.17), and SW and MW had a moderate difference in pulse rate at 2.15 pulses/second (95%CI = 1.21–3.10). Conversely, we found no difference in pulse number among lineages (*p*=0.426; SW = 15.20, MW = 15.13, NE = 14.99). Visual inspection of population averages show little association of pulse number with geography (Supp. Fig. 8), but there is a clear association between faster pulse rate and the SW lineage range, with faster calls in MW *H. versicolor* occurring closer to the MW/SW contact zone (Fig. 3a).

NMM analysis of all *H. versicolor* calls support a single component as the best model with BIC = 566.16 and a model of two components with BIC = 551.88 (Supp. Fig. 9). NMMs fit to components defined by the mitochondrial lineages show high precision for separating NE and SW lineages, but categorizes most MW calls as belonging to the NE lineage (Supp. Fig. 10). Analyses where two lineages were collapsed show high precision when NE and SW lineages are separated and fails to properly separate MW calls as a separate component when NE and SW were combined. The collapsed lineage model with the highest BIC and lowest classification error (CE) was the model where SW was separate from NE and MW with BIC = 342.77 and CE = 0.079, the second highest was the model where NE was separate from SW and MW with BIC = 341.57 and CE = 0.174. The SW and NE collapsed model had the lowest BIC with BIC = 331.94 and CE = 0.172 (the low classification error here driven by accurately classifying all 251 SW + NE outweighing the misclassification of the 52 MW individuals). Although the NMM of all calls suggested a single component, the results of the lineage specific NMM suggest the SW lineage to be the most distinct, corroborating the results of our randomization test.

### Acoustic Analysis of H. chrysoscelis

Our analysis of phenotypic evolution using PhylogeneticEM suggest that acoustic signals of *H. chrysoscelis* are evolving under three different selective regimes with respect to pulse rate and pulse number (Fig. 3c). Broadly, a low pulse rate and high pulse number in the eastern part of the species’ range and a high pulse rate and low pulse number in the west characterize these selective regimes (Fig. 3b-d; Supp. Fig. 11). Looking more closely, it appears that the selective regimes of the Eastern/Central lineages are split across the Appalachian Mountains (Fig. 3b; Fig. 3d; Supp. Fig. 11). This result is consistent with the pattern of northward migration suggested by our biogeographic analyses, and it conforms to the estimated population structure in this area (Booker et al. 2022). Importantly, although no previous work has demonstrated a relationship between geography nor genealogy and pulse number (e.g., Supp. Fig. 11), our analysis demonstrates an association between pulse number and phylogenetic relationships in *H. chrysoscelis* (Fig. 3c). Finally, an important caveat of these results is that PhylogeneticEM assumes there is no gene flow across the phylogeny. Given previous results (Booker et al. 2022), we know this assumption is violated. However, we do believe that the pattern shown is clear and in agreement with other analyses here and from previous work despite this violation.

NMM analysis of all *H. chrysoscelis* individuals suggests a model with two components as the best model with a BIC = 1628.41, and one and three component models had BIC = 1539.99 and BIC = 1613.10 respectively (Supp. Fig. 12). The geographic distribution of these components largely split the calls between east and west populations by pulse rate (Fig. 4b), similar to the PhylogeneticEM analysis. At a finer scale, again similar to the PhylogeneticEM analysis, the Eastern and Central lineage populations are represented by the two components separately, divided by the Appalachian Mountains. An NMM fit to components defined by the 3 regimes split by PhylogeneticEM showed high precision for Eastern (green in figure Fig. 4a) and Western (black in figure Fig. 4a) individuals, but classified Central (gray in Fig. 4a) as 44% and 33% Western and Eastern respectively (Supp. Fig. 13). NMMs where two regimes are collapsed showed generally high precision except for the model where Eastern and Western regimes were collapsed into each other, and this model had the lowest BIC and CE at BIC = 570.98 and CE = 0.23. The model where the Western regime was separate from Eastern and Central had the highest BIC but second lowest CE at BIC = 639.56 and CE = 0.145. The model where the Eastern regime remained separate from Western and Central had the second highest BIC and lowest CE at BIC = 629.25 and CE = 0.100. Ultimately, these results laregely agree with the PhylogeneticEM results, suggesting two distinct distributions of calls with the Appalachian Mountains playing a significant role in driving call differences of Eastern/Central *H. chrysoscelis*.

**Figure 4.**
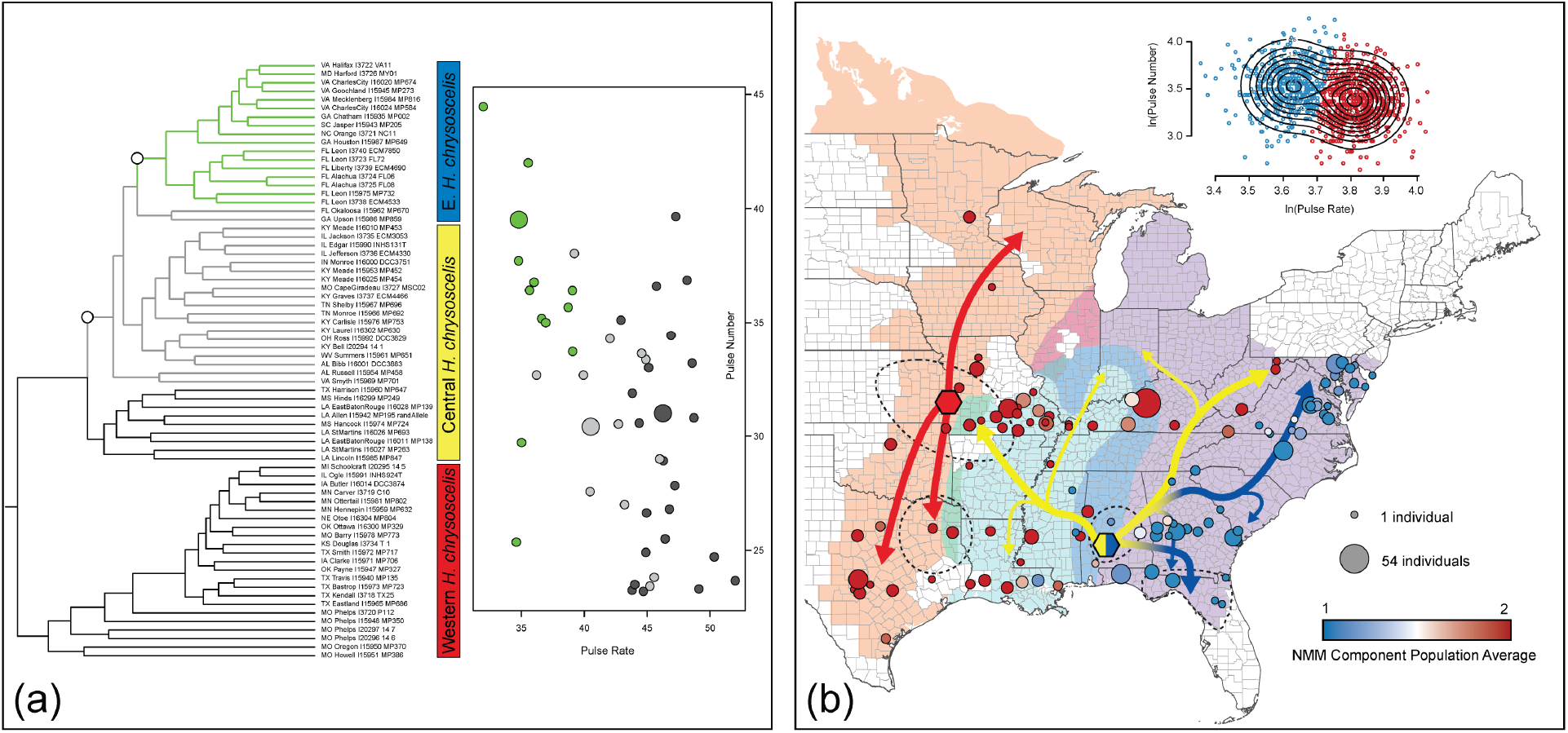
Acoustic evolution of *H. chrysoscelis* in the context of biogeography. (a) Phylogeny and acoustic regimes estimated from PhylogeneticEM analysis. On the phylogeny, white circles depict regime shifts at a given node, and branches are colored with respect to their selective regime. Circle colors correspond to the regime colors on the phylogeny, and small circles are the observed pulse rate and pulse number means for a population, while large circles represent the optimum value of each regime estimated by PhylogeneticEM. Western (red), Central (yellow), and Eastern (blue) *H. chrysoscelis* are lineages delimited in Booker et al. (2022). (b) Biogeography and acoustic evolution of *H. chrysoscelis*. Background colors show the estimated ranges of mitochondrial lineages from Fig. 1. Colored circles represent the average component of individuals in each population from the NMM analysis, and the size of circles corresponds to the number of individuals analyzed in each population. Inset figure shows the distribution of components as it relates to pulse rate and pulse number, with the density of points in the components overlayed. Colored hexagons depict the ML estimated ancestor location for Western (red) and Eastern/Central (blue/yellow) lineages from panel a. Colored arrows show the expansion of lineages out of the ancestor location summarized from Supp. Fig. 6 and Supp. Fig. 7, with colors corresponding to present day Western (red), Eastern (blue), and Central (yellow) lineages from panel a. Dashed areas highlight proposed refugia that have been used by *H. chrysoscelis*.

## Discussion

The results from this study and previous estimates demonstrate the relationship between glaciation cycles, migration, and species interactions in North American gray treefrogs and highlight the effect of a complex biogeographic history on acoustic evolution in this group. In *H. versicolor*, we found evidence that both the original formation of *H. versicolor*, as well as one of two mitochondrial introgression events, are associated with the limit of the Laurentide ice sheet during the Quaternary glaciations approximately when those events occurred (Table 1; Limits of the Laurentide ice sheet during Illinoian and Wisconsin glacial limits shown in Fig. 1a,b). For the second mitochondrial introgression event, we see a marked difference of call phenotypes in the SW *H. versicolor* lineage (Fig. 3a) that is correlated with the faster pulse rate calls seen in CSW *H. chrysoscelis* (Fig. 3b) with whom they share a mitochondrial lineage. Within *H. chrysoscelis*, we found two largely distinctive call types—a slow pulse rate and high pulse number call in eastern populations, and a faster pulse rate but lower pulse number call in western populations (Fig. 3b; Fig. 4a). In general, call diversity in this species shows a strong association with nuclear phylogenetic relationships. Furthermore, the totality of evidence from this study and Booker et al. (2022) suggest the east-west split in calls evolved prior to the expansion of *H. chrysoscelis* out of east and west refugia, and that additional geographic isolation and east-west gene flow has contributed to the wide diversity of calls observed throughout the species’ range.

### Biogeographic History of H. versicolor

An interesting finding of this study is the association of major evolutionary events with the range limit of the Laurentide ice sheet approximately around when these events would have occurred (Table 1; Booker et al. 2022; Fullerton 1986; Curry et al. 2011). A significant hurdle to polyploid establishment is the relative shortage of suitable mates when polyploids are first formed (i.e. minority cytotype exclusion, Levin 1975). One way that established polyploids may have overcome this issue is by successfully expanding into unoccupied habitat either by shifting ecological niches or by forming on the range periphery of the progenitor species (Stebbins 1985; Brochmann et al. 2004; Parisod et al. 2010; Van de Peer et al. 2017). One prediction of this hypothesis is that we should observe a positive association with overall polyploid species diversity and the proximity of glaciated areas (Brochmann et al. 2004; Novikova et al. 2018). Indeed, a recent review of the geographic diversity of plant polyploids showed a positive relationship between the extent of de-glaciation and polyploid abundance (Rice et al. 2019), and a recent review on animal polyploids demonstrated glaciation as the top predictor of discrepancy between polyploid and diploid diversity, with polyploids more likely to occur in previously glaciated environments (David 2022). Our results demonstrating the origin of *H. versicolor* (i.e. NE *H. versicolor*) near the limit of the Laurentide ice sheet in the Northeastern U.S. are consistent with these observations. Given the timing estimates of WGD are all during recent Quaternary glaciation cycles, it is possible that de-glaciation during this period would have provided the necessary conditions for *H. versicolor* to become established.

Our results suggest that the origin of MW *H. versicolor* is also associated with Quaternary glaciations (Fig. 2b). Work from Booker et al. (2022) suggests this lineage was formed when NE *H. versicolor* hybridized with a now extinct population of *H. chrysoscelis*, rather than as an independently formed polyploid as suggested by Holloway et al. (2006). Under this model, MW *H. versicolor* necessarily formed after NE *H. versicolor*, and given the timing of whole genome duplication inferred from ABC (100Ka, Booker et al. 2022), it is most likely that the MW lineage formed after the Illinoian glacial period (approximately 190–130 Ka Curry et al. 2011) but prior to the Wisconsin glacial period (approximately 60–12.5 Ka Curry et al. 2011). Under this scenario, the growth of the Laurentide ice sheet leading into the Wisconsin glacial period would have separated *H. versicolor* populations that had expanded out of New England. Following isolation, *H. versicolor* around central Illinois would have hybridized with local *H. chrysoscelis*, integrating their mitochondrial genomes before expanding back east and making secondary contact with NE *H. versicolor* after the Last Glacial Maximum (LGM). However, given the coalescent timing from previous mitochondrial phylogenetic analyses 1, there is uncertainty around the precision of this estimate. Nonetheless, regardless of the estimate used, the timing of these events is approximately near periods of glacial cycles in the late Quaternary.

Although the origin of SW *H. versicolor* is not directly on the edge of a Quaternary glacial limit, its estimated location in the Arkansas river valley between the Ouachita and Boston mountain ranges is within previously proposed Pleistocene refugia (Delcourt and Delcourt 1984; McLachlan et al. 2005; Soltis et al. 2006; Fontanella et al. 2008). Perhaps most relevant to gray treefrogs, Austin et al. (2002; 2004) proposed this area as a refugium for *Pseudacris crucifer*, another Nearctic Hylid whose current range largely overlaps that of *H. versicolor* and *H. chrysoscelis*, and like gray treefrogs, spans a wide array of ecosystems from the Southeastern Coastal Plain to the Arctic Circle. Given the recent formation (Table 1), mitochondrial introgression (Booker et al. 2022), and the distinct call in this lineage (Fig. 3a), future work should further investigate the relative contribution of glaciation cycles to the evolution of SW *H. versicolor* (e.g., reducing population sizes or driving species interactions). The distinctive patterns observed in this lineage suggest this work could provide unique insights into how abiotic factors influence the diversification of polyploids.

Importantly, the scenarios presented above, though plausible in light of our understanding of the complex, assume that the single origin model is correct. Though this model was supported by several lines of evidence, some of its details remain unclear. Booker et al. (2022) found conflicting evidence as to which *H. versicolor* lineage was a progenitor of SW *H. versicolor*, though the majority of evidence suggests a MW *H. versicolor* progenitor. Ambiguity also exists as to the number of mitochondrial introgression events in SW *H. versicolor*. Here, we provide some clarity to the SW lineage’s history, as the phylogenies of each species individually used here showed a greater association between geography and phylogenetic relationships in this lineage than did the multi-species phylogeny from Booker et al. (2022). Future efforts should focus on further clarifying the relationships among lineages and using more intricate demographic models to explicitly test the hypotheses presented.

### Biogeographic History of H. chrysoscelis

Our biogeographic analyses revealed several probable refugia that were utilized by *H. chrysoscelis* and demonstrate how migration out of these refugia has influenced the current population structure of this species. Evidence for an east-west split in *H. chrysoscelis* has been available for nearly half a century (Gerhardt 1974; Ralin 1976). Work from Booker et al. (2022) further separated Eastern *H. chrysoscelis* into two lineages (Eastern and Central). Despite this structure, gene flow between the lineages appears to occur wherever they interact. In the present study, we estimated the geographic origin of the east-west split in *H. chrysoscelis* and further estimated the migratory expansion routes out of proposed glacial refugia.

We estimated the East Gulf Coastal Plain in southern Alabama as the center of origin of Eastern and Central *H. chrysoscelis* (Fig. 1e). This location has repeatedly been identified as a probable refugium across multiple taxonomic groups (Swenson and Howard 2005; Soltis et al. 2006), and it appears to have been a particularly important refugium for amphibians (Rissler and Smith 2010). Our results and the previously estimated coalescent times (Table 1) suggest this location, and the broader Southeastern Coastal Plain and Piedmont in general, have persisted as stable habitat for *H. chrysoscelis*—providing a source for recolonization of eastern North America. The coalescent time of all *H. chrysoscelis* was estimated at 1.26 Ma, and the coalescent time of all Eastern and Central *H. chrysoscelis* was estimated at 811 Ka (Fig. 1; Table 1). These time estimates, alongside the significantly lower dispersal rate of *H. chrysoscelis* in this area that didn’t expand northward towards the Appalachian Mountains, suggest that *H. chrysoscelis* in the Southeast have remained in this area through multiple glaciation cycles. We also found evidence from both our biogeographic and acoustic analyses that North Florida was an additional refugium for *H. chrysoscelis*, and that this area holds additional diversity from refugial populations (see Acoustic Signal Evolution in Diploid *H. chrysoscelis*). Support for this area as a refugium also exists in the literature. This area holds relict populations of several species of plants (Parks et al. 1994; Soltis et al. 2006), was previously identified as a refugium for other ectotherms (e.g. *Heteranria formosa* Bagley et al. 2013), and is home to a high number of endemic species (Harper 1914; Neill 1957; Means 1977; Means and Krysko 2001; Avise 2000).

Though Western *H. chrysoscelis* have persisted through multiple glaciation cycles, the estimated coalescent timing of extant Western *H. chrysoscelis* is much more recent than the timing for Eastern and Central *H. chrysoscelis*. Booker et al. (2022) demonstrated coalescence of extant Western *H chrysoscelis* individuals approximately 262 Ka (Fig. 1; Table 1), and our analysis in the present study suggests this event occurred near the intersection of Missouri, Arkansas, Oklahoma, and Kansas. This broad area is within the Ozarks and Great Plain regions which have been previously proposed as refugia for multiple species including anurans (McLachlan et al. 2005; Soltis et al. 2006). We identified evidence for one other refugium in eastern Texas, where Western *H. chrysoscelis* may have persisted during the Wisconsin glacial period. The same refugium, which has been suggested for multiple taxa (Swenson and Howard 2005), including other treefrogs (Lemmon et al. 2007a; Barrow et al. 2015) and rat snakes (Burbrink 2002), may have been of particular importance because much of the Midwest was significantly more arid during interglacial periods than it is currently (Bartlein et al. 1998; Lemmon et al. 2007b). Importantly, previous analyses suggest the existence of two more extinct lineages of *H. chrysoscelis*, whose mitochondrial lineages remain preserved in NE and MW *H. versicolor* (Booker et al. 2022). Though the cause of their extinction is unknown, based on the range of their respective polyploid relatives these lineages most likely existed farther north than the extant lineages of *H. chrysoscelis* and likely utilized additional refugia.

### Acoustic Signal Evolution in Tetraploid H. versicolor

The results from our analyses demonstrate a lack of pulse number variation across *H. versicolor* lineages, but significant variation in pulse rate across lineages (Fig. 3b). The most significant result from our acoustic analysis in *H. versicolor* was the association of faster pulse rate calls with the SW mitochondrial lineage. Although the “genomic shock” (McClintock 1984) of polyploidy likely had little effect on most phenotypic characters of *H. versicolor* due to its autopolyploid origin, polyploidization did have a substantial effect on their physiology (Ueda 1993; Keller and Gerhardt 2001; Tucker and Gerhardt 2012), and consequently, the vocalizations of this species. A significant implication from our observations is that calls in polyploids may shift rapidly following novel interactions with diploid relatives, and that these shifted calls can persist in a population as it expands its range (Fig. 3a). Calls in this SW *H. versicolor* have significantly faster pulse rates than other *H. versicolor*, and the differences observed in this lineage have a nearly identical directional similarity in the percent difference in pulse rate of *H. chrysoscelis* in the same area. Given that a shift in PR towards local *H. chrysoscelis* calls would likely result in a greater proportion of costly diploid-tetraploid hybridizations, exactly why the unique SW call has proliferated needs to be considered.

Based on their association, a logical consideration is the effect mitochondrial introgression could have on *H. versicolor* calls. Mitochondrial introgression has a unique and disproportionate effect on evolution due to the intimate interaction between nuclear and mitochondrial coded genes that are necessary for essential cellular functions (Levin 2003; Burton et al. 2013; Sloan et al. 2017). As Burton et al. (2013) suggest, the establishment of successful cytonuclear interactions is critical to the success of hybrid lineages. This effect is further exaggerated in polyploids (Sharbrough et al. 2017), and organelle-target genes are among the first to return to single copy following WGD (De Smet et al. 2013). The most recent common ancestors of *H. versicolor* mitochondria are the oldest in the whole gray treefrog complex (SW:MW 1.6 Ma, NE:(SW:MW) 1.8 Ma) yet the initial WGD and all subsequent introgressions occurred within the last 262,000 years (Table 1). These dates suggest SW *H. versicolor* nuclear alleles that are not the result of recent gene flow evolved in concert with other mitochondria for approximately 1.3 million years. This disparity, and the frequency of diploid-tetraploid gene flow alongside the lack of contemporary observations of mitochondrial introgression, suggest cytonuclear interactions may play a role in this complex. If this were the case, in SW *H. versicolor* the introgression of CSW *H. chrysoscelis* mitochondria then may have had a disproportionate effect in driving the observed call divergence, because selection would have favored alleles of organelle-targeted genes that co-evolved with the CSW mitochondria—resulting in the over-representation of local *H. chrysoscelis* alleles in SW *H. versicolor*. Curiously, we do not see the same pattern in MW *H. versicolor* that overlap with faster pulse rate *H. chrysoscelis* except where they also interact with SW *H. versicolor* (Fig. 3a), even though diploid to tetraploid gene flow has likely been a continual occurrence in MW *H. versicolor* since its formation (Booker et al. 2022). A detailed look at the disparity between SW and MW calls despite similar evolutionary histories would be a fruitful avenue for future study, and may provide insights into the genomic components of acoustic communication.

### Acoustic Signal Evolution in Diploid H. chrysoscelis

Results from this study demonstrate that *H. chrysoscelis* acoustic signals are evolving under two to three different selective regimes specific to geographic location and phylogenetic clades (Fig. 3b-d; Fig. 4; Supp. Fig. 11), and that call evolution is related to their biogeographic history. The three defined nuclear genetic lineages are largely differentiated by a low pulse rate and high pulse number in the Eastern lineage, a high pulse rate and moderate pulse number in the Central lineage, and a high pulse rate and low pulse number in the Western lineage. The exception, likely due to gene flow, is Central lineage individuals in Alabama, Georgia, Florida, and Mississippi, which have a slower pulse rate more similar to the Eastern lineage. Interestingly, the observed nuclear and phenotypic patterns are not phylogenetically concordant: the Central lineage is genetically more similar to the Eastern lineage, but generally has a pulse rate most similar to the Western lineage.

Our biogeographic analysis suggests that much of the variation in Eastern/Central *H. chrysoscelis* calls may be linked to the expansion of refugial populations that evolved different acoustic properties. In particular, the more recent expansion of *H. chrysoscelis* north in the eastern part of its range, with unique lineages expanding east and west of the Appalachian Mountains, may have resulted in the divergence of calls between these two lineages despite sharing a recent common ancestor and their close geographic proximity. Whereas *H. chrysoscelis* on the western slope of the Appalachian Mountains have a faster pulse rate characteristic of Western *H. chrysoscelis*, populations on the east side have a much slower pulse rate. Results from our comparative analyses suggest a more recent evolution of this slower pulse rate, which is shared across southeast, east coast, and east Appalachian populations. Exactly when and where the slow-pulse rate phenotype evolved is unclear. We find the slowest pulse rates in eastern Florida and Georgia, and calls from populations in the Big Bend region and peninsular Florida demonstrate slowest pulse rate and highest pulse number in all of *H. chrysoscelis* (Fig. 3b, Supp. Fig. 11). In the present study, we identified several lines of evidence that suggested this area as a stable refugium for *H. chrysoscelis*. Therefore, an Appalachian origin of this call seems unlikely. A more likely scenario is that this call evolved in southeast populations, and stable Florida populations may have helped this call phenotype to persist through Quaternary climatic oscillations.

Unlike Eastern/Central lineages, we find limited evidence for phenotypic shifts in Western *H. chrysoscelis* as a result of expansion—suggesting that calls in this lineage have undergone limited diversification following the lineage’s initial isolation. Western *H. chrysoscelis* lack a similar physical barrier to the Appalachians that separate call types in Eastern/Central *H. versicolor*, and it is likely that the relative elevational homogeneity of the Western lineage’s range is a cause of this disparity.

## Conclusions

In the present work we examined the biogeographic history of the North American gray treefrog complex and assessed the history of call evolution in this context. The details herein outline the supposed paths that various lineages of this group took throughout the Quaternary that led to its present day distribution and acoustic phenotypic diversity. Still, the precise forces which acted to carve out this path, and excluded others, remain unclear. Why, if these species exist in sympatry over much of their range, are they not able to coexist elsewhere? Or perhaps more accurately, given their extreme similarity, how are these species able to coexist over much of their range and in such a diversity of habitats? To the extent that evolution is predictable, the answer may lie in the specific lineages that came into contact and in the forces that shaped those lineages—perhaps in the specific adaptations in those areas, the acoustic properties of the lineages when they were formed and interacted, or in the genetic consequences that shaped the trajectory of expanding and contracting populations. Alternatively, the answer may lie in the habitats themselves, the qualities of which have been shaped by repetitive ecological turnover and drastic climatic oscillations. Though we cannot replay life’s tape to uncover the answers to these questions, we hope the work presented here, alongside previous work in the complex, provides some basis for which these lines of inquiry can continue to be addressed.

## Acknowledgements

We would like to thank Allison Welch, Joe Mitchell, and Jen Henderson for providing acoustic data; Hank Bass, Jim Bogart, David Hillis, Joseph Travis, and Rafael Marcondes for helpful feedback during project development and analysis. This work was supported by the National Science Foundation through an NSF GRF (DGE 1449440) and NSF DEB (1120516), and by the University of Missouri Research Council (URC-15-106).

## Data Accessibility

All data used for this study are openly available in Dryad at https://doi.org/10.5061/dryad.z8w9ghxhr.

## Supplementary Information

**Supplemental Figure 1.**
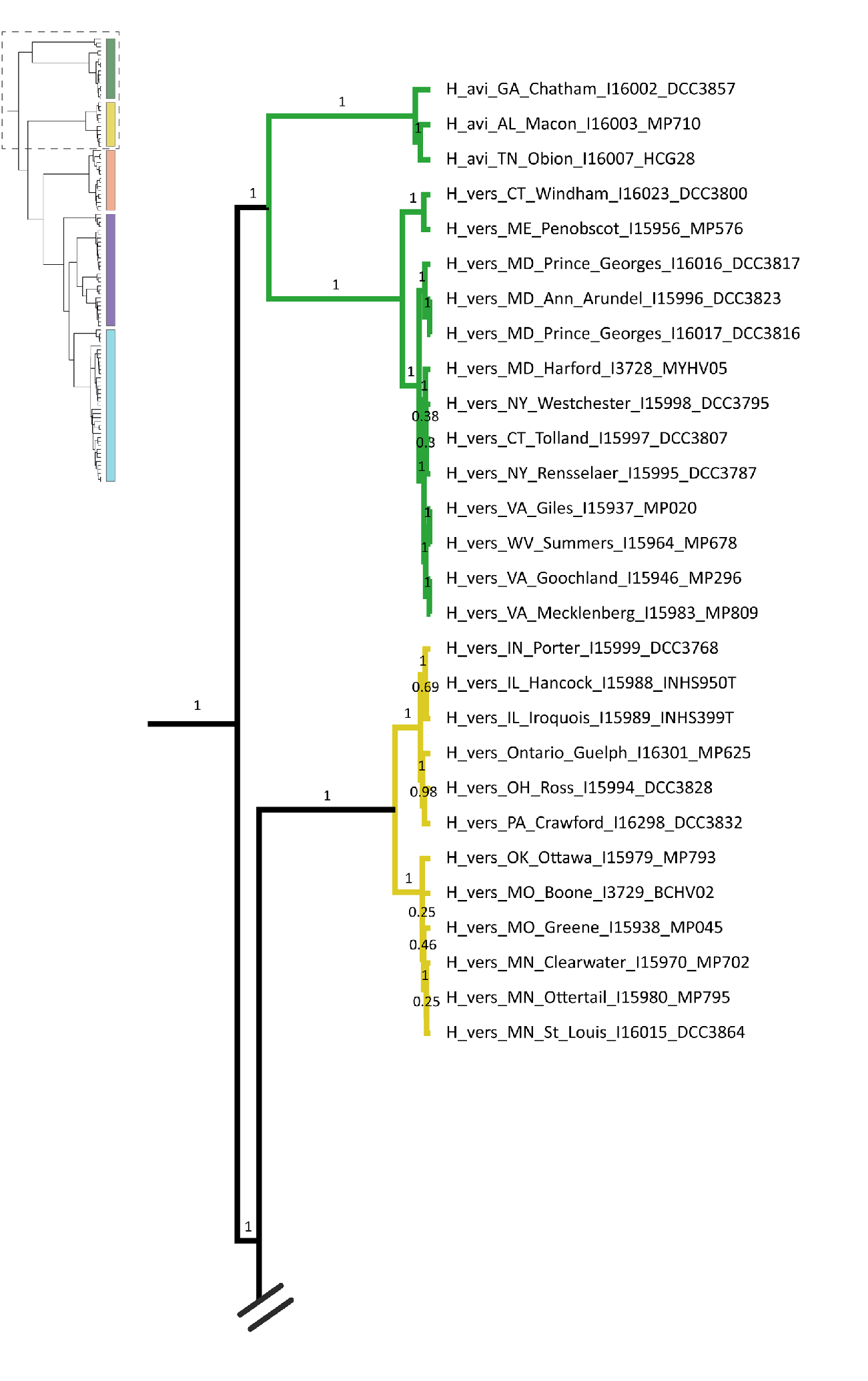
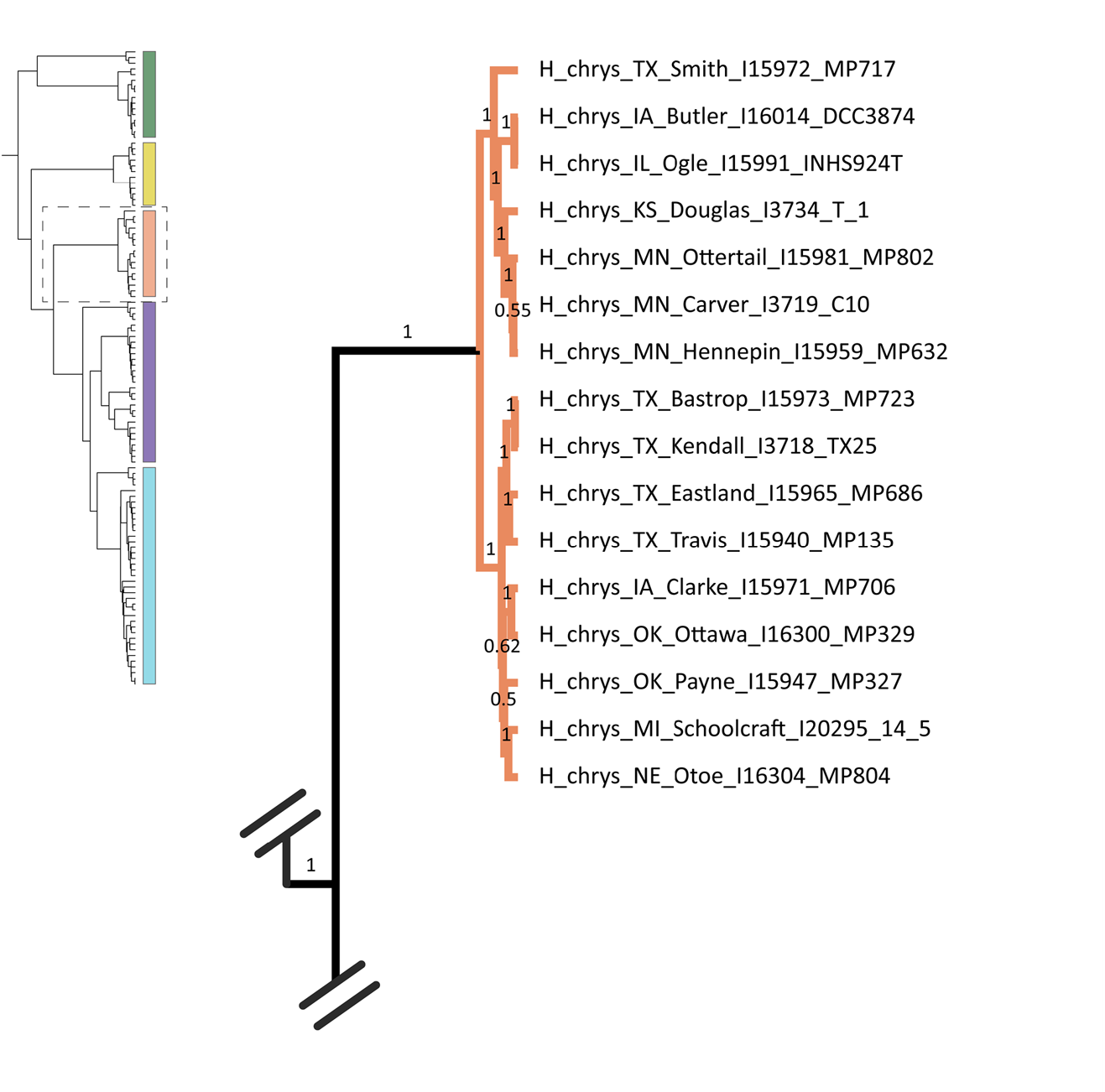
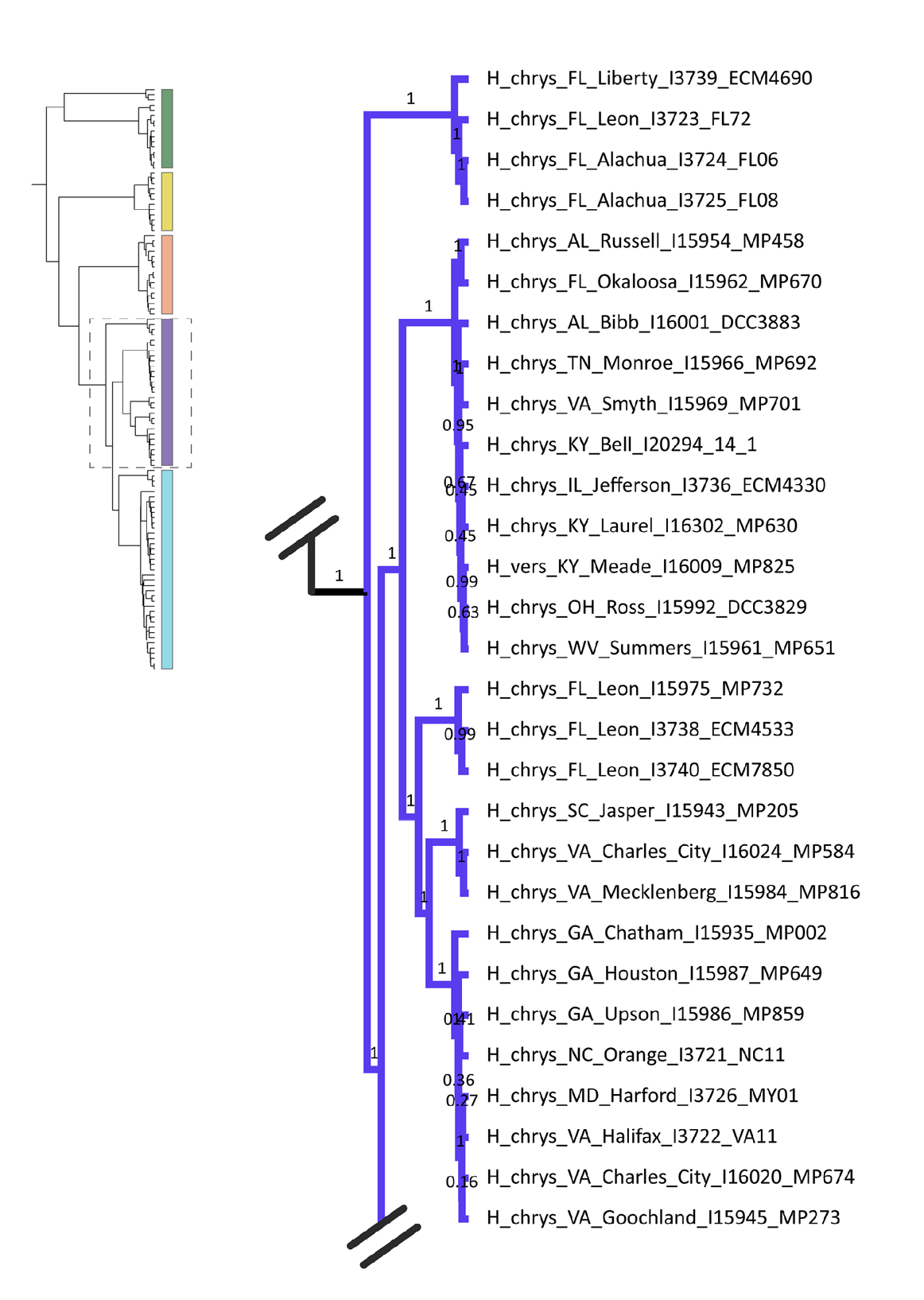
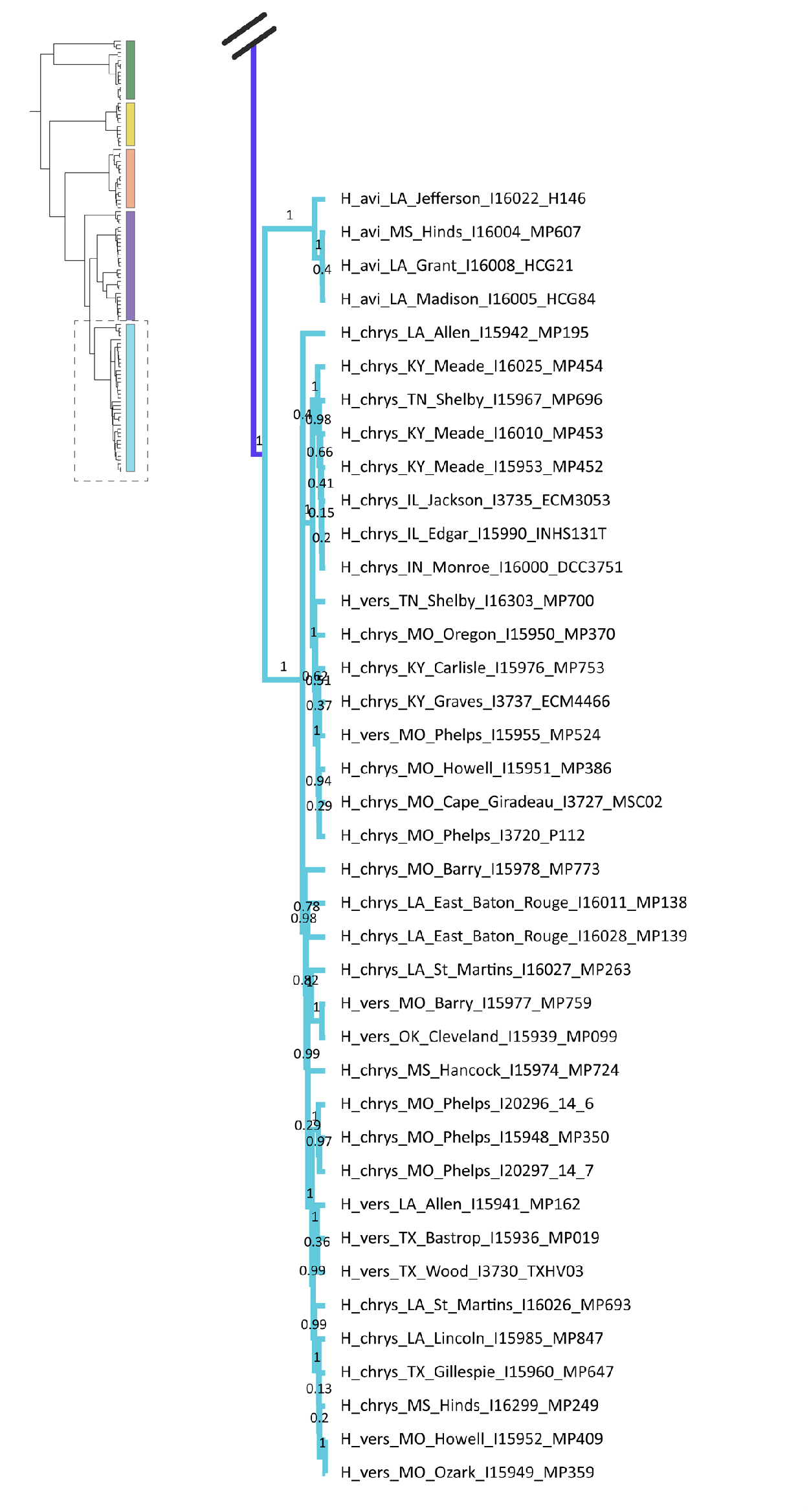
Expanded mitochondrial tree estimated for *Hyla avivoca* (H avi), *Hyla chrysoscelis* (H chrys), and *Hyla versicolor* (H vers). shown in Fig. 1 from Booker et al. (2022). Species names are followed by state, county, internal ID, and museum ID. Branch labels show posterior probability. Clade colors represent delineated mitochondrial lineages.

**Supplemental Figure 2.**
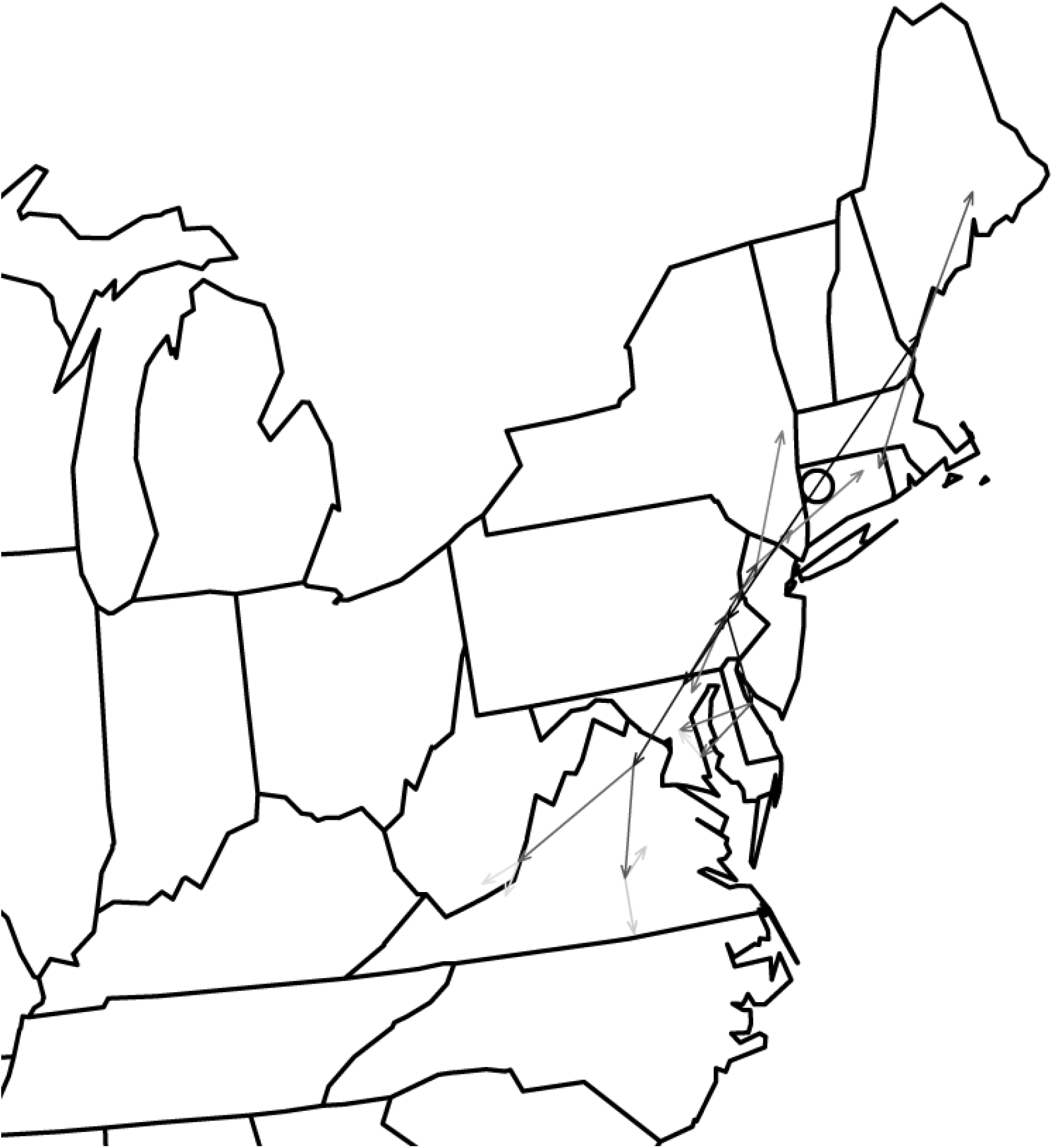
Full expansion map of NE *H. versicolor* from Phylomapper analyses using a tree generated from whole mitochondrial genomes. The open circle represents the ancestral location of all NE *H. versicolor*. Arrows are colored with respect to phylogenetic depth, with darker arrows originating from deeper nodes and lighter arrows originating from more shallow nodes on the tree.

**Supplemental Figure 3.**
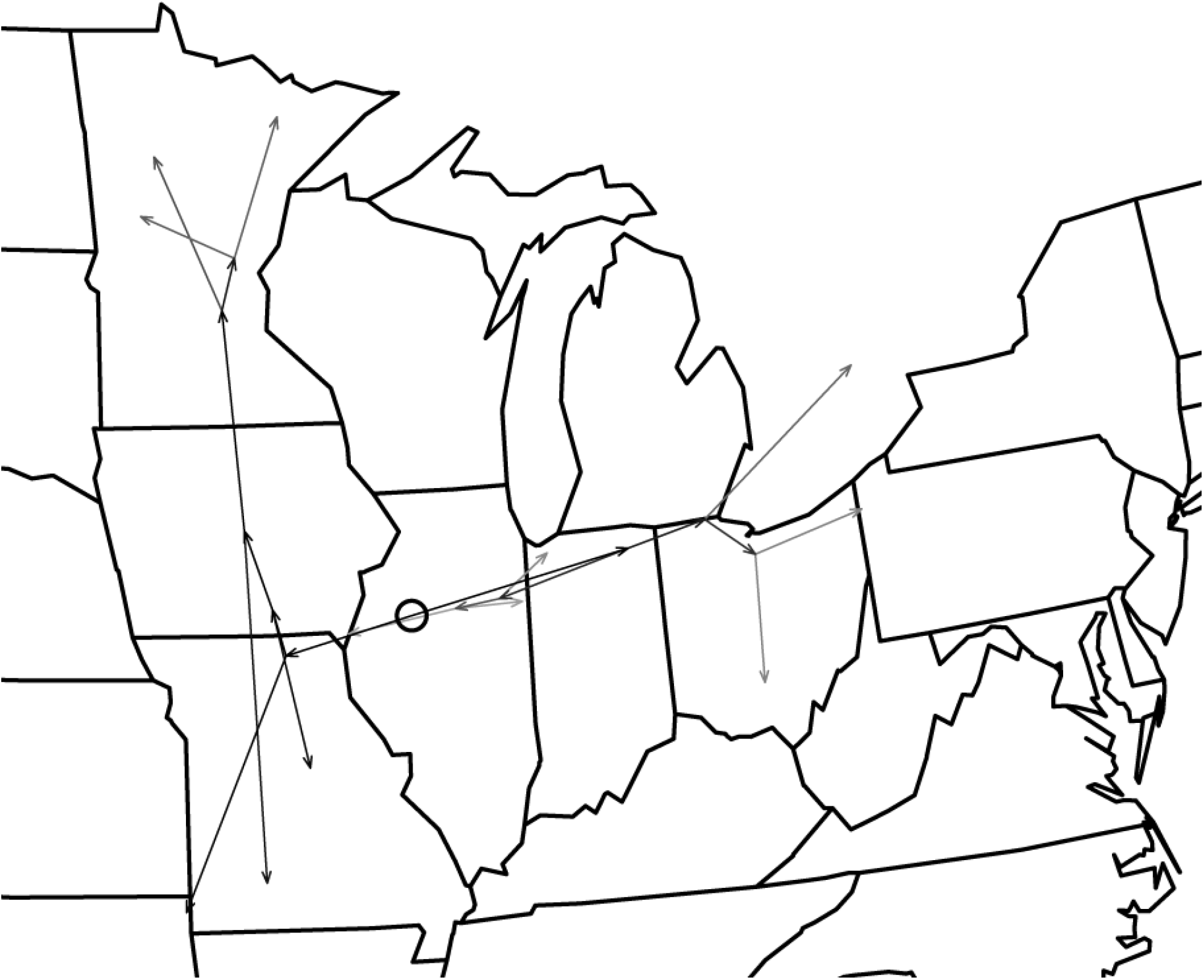
Full expansion map of MW *H. versicolor* from Phylomapper analyses using a tree generated from whole mitochondrial genomes. The open circle represents the ancestral location of all MW *H. versicolor*. Arrows are colored with respect to phylogenetic depth, with darker arrows originating from deeper nodes and lighter arrows originating from more shallow nodes on the tree.

**Supplemental Figure 4.**
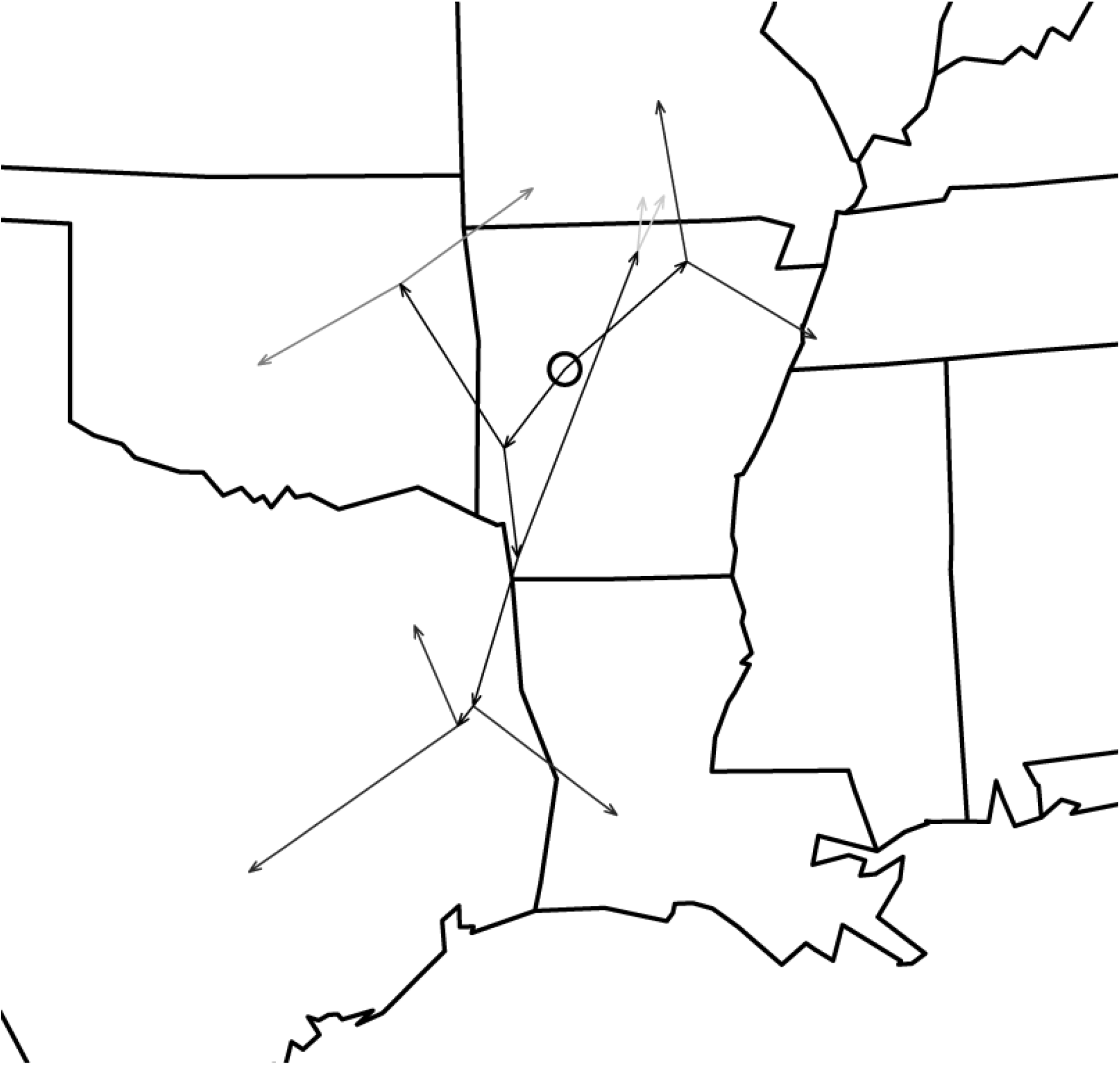
Full expansion map of SW *H. versicolor* from Phylomapper analyses using a tree generated from whole mitochondrial genomes. The open circle represents the ancestral location of all SW *H. versicolor*. Arrows are colored with respect to phylogenetic depth, with darker arrows originating from deeper nodes and lighter arrows originating from more shallow nodes on the tree.

**Supplemental Figure 5.**
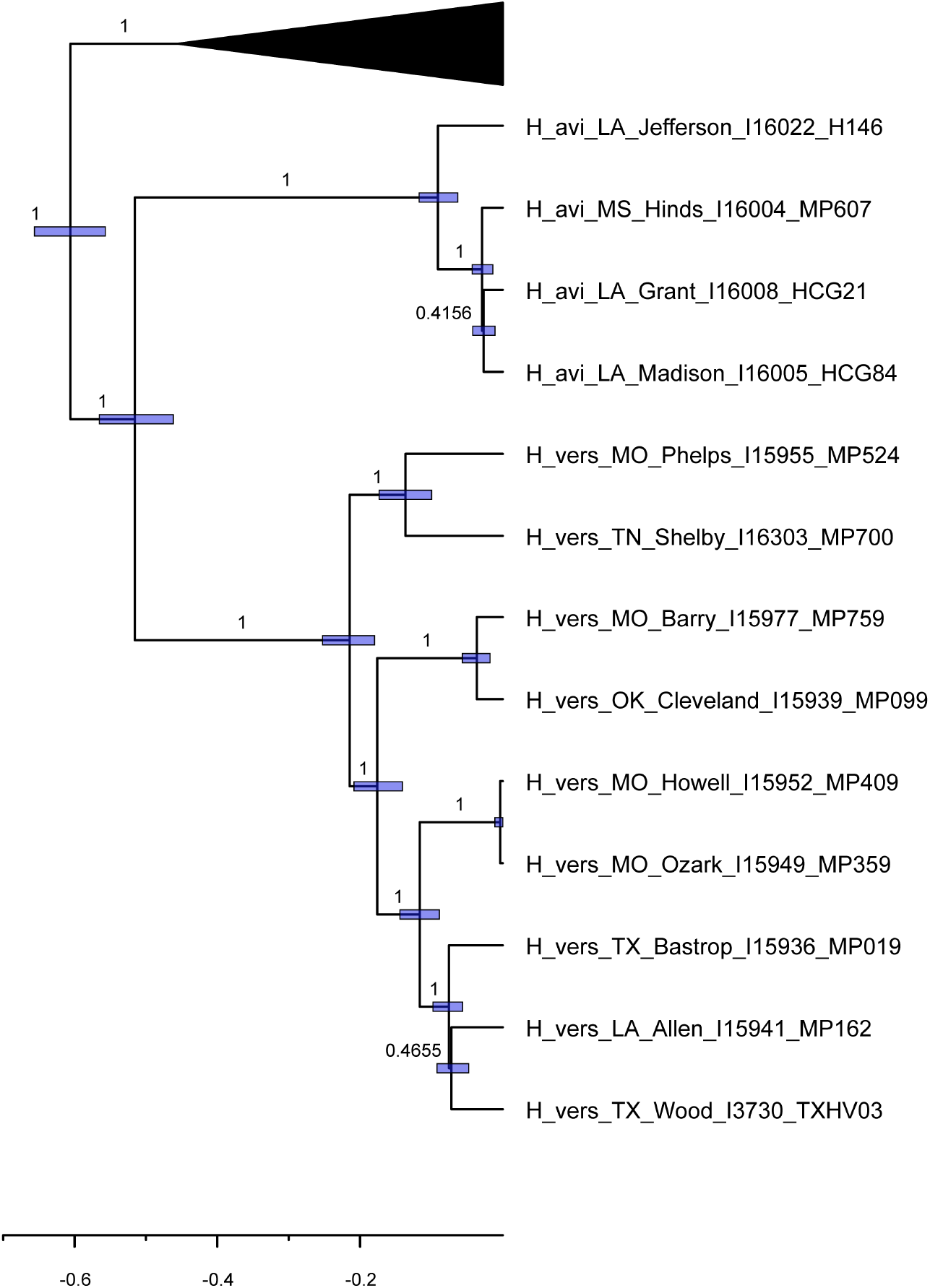
Mitochondrial genome tree of SW *H. versicolor* after removing CSW *H. chrysoscelis*. Branch labels represent posterior probability. Bars represent 95%CI of coalescent time in millions of years.

**Supplemental Figure 6.**
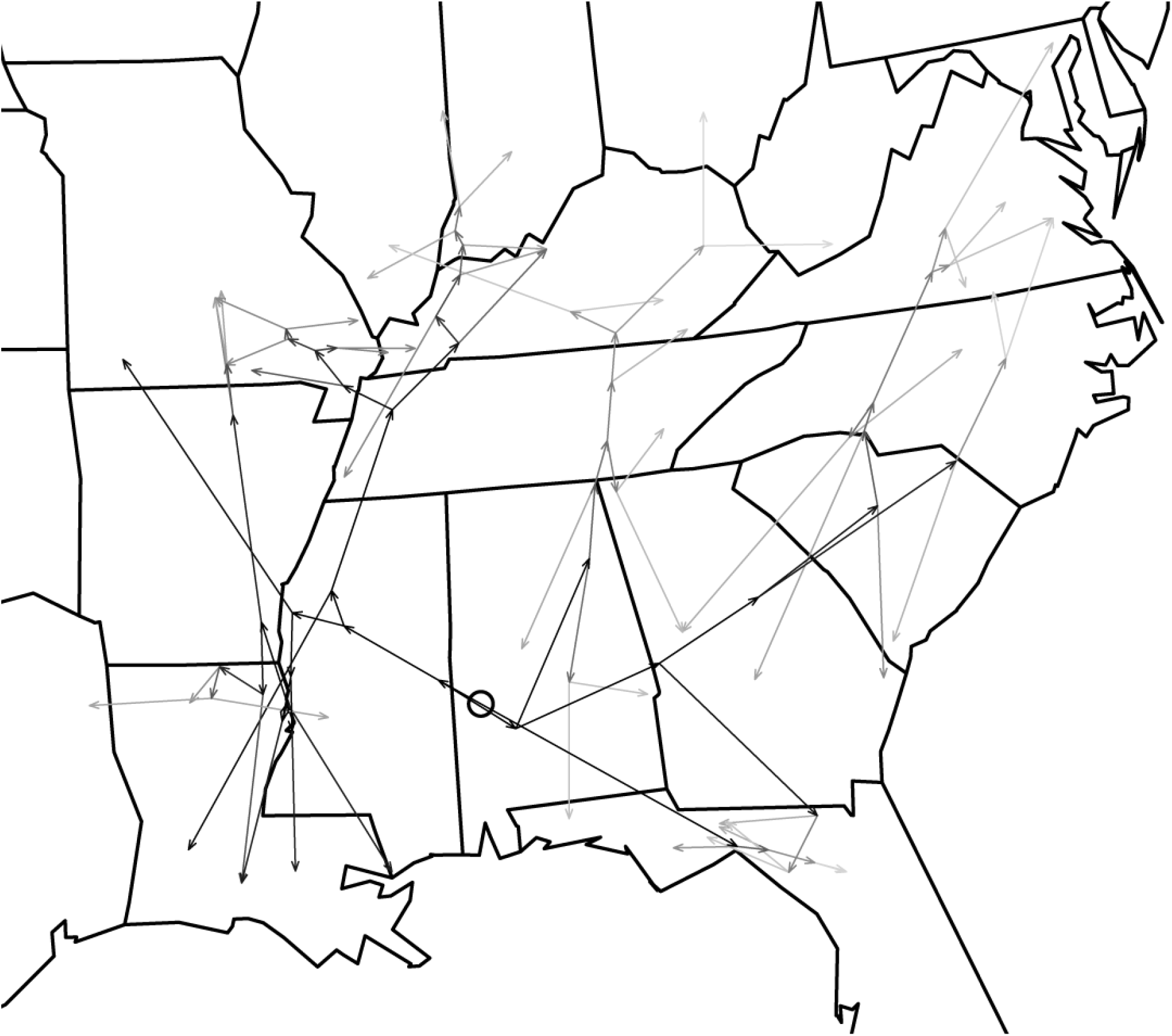
Full expansion map of Eastern and Central *H. chrysoscelis* from Phylomapper analyses using a tree generated from whole mitochondrial genomes. The open circle represents the ancestral location of all Eastern and Central *H. chrysoscelis*. Arrows are colored with respect to phylogenetic depth, with darker arrows originating from deeper nodes and lighter arrows originating from more shallow nodes on the tree.

**Supplemental Figure 7.**
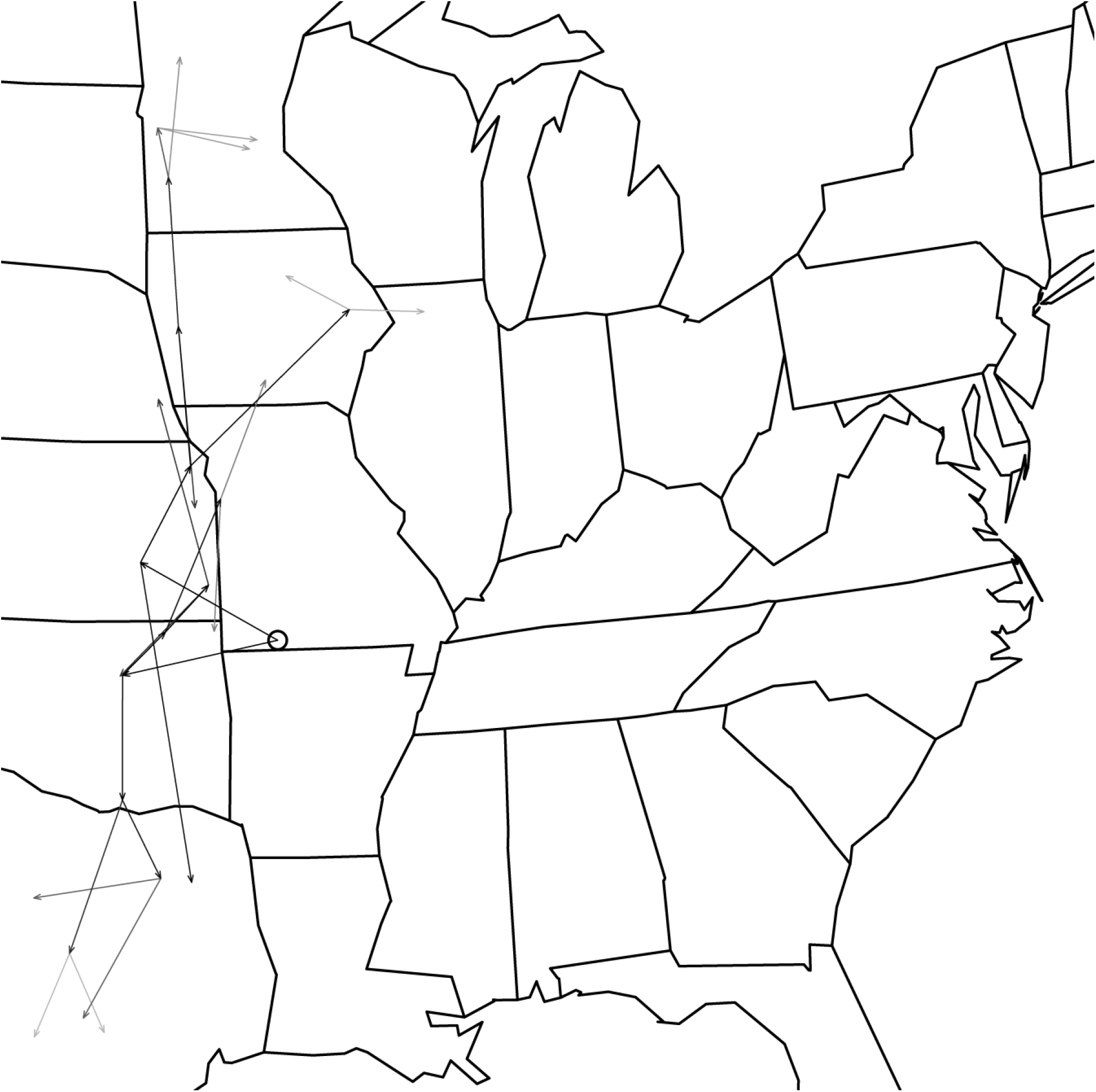
Full expansion map of Western *H. chrysoscelis* from Phylomapper analyses using a tree generated from whole mitochondrial genomes. The open circle represents the ancestral location of Western *H. chrysoscelis*. Arrows are colored with respect to phylogenetic depth, with darker arrows originating from deeper nodes and lighter arrows originating from more shallow nodes on the tree.

**Supplemental Figure 8.**
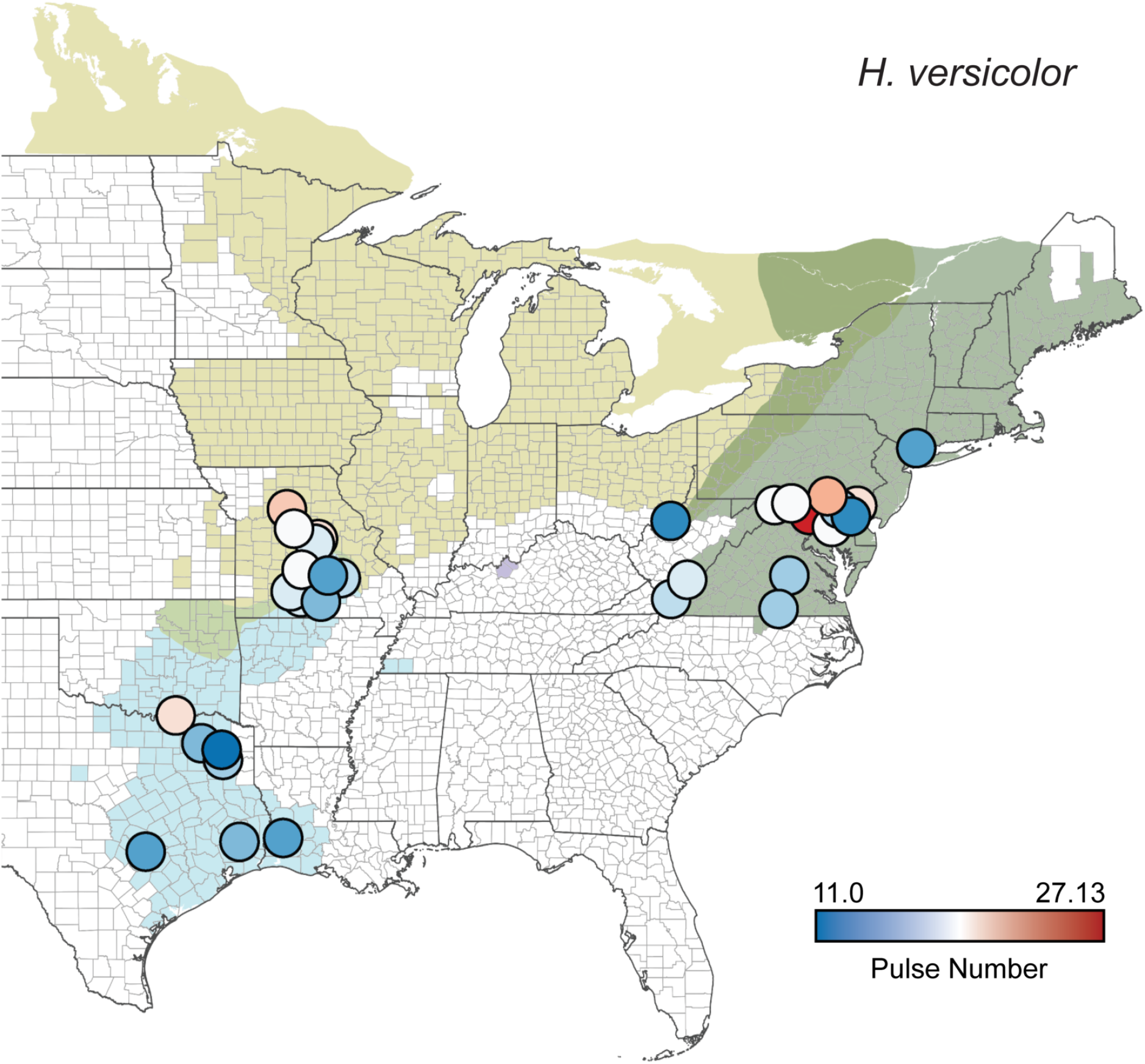
Call pulse number (total number of pulses in a call) in *H. versicolor*. Circles represent averages for a single population. Points are colored from slow pulse number (blue) to high pulse number (red). Background colors correspond to mitochondrial lineages from Fig. 1.

**Supplemental Figure 9.**
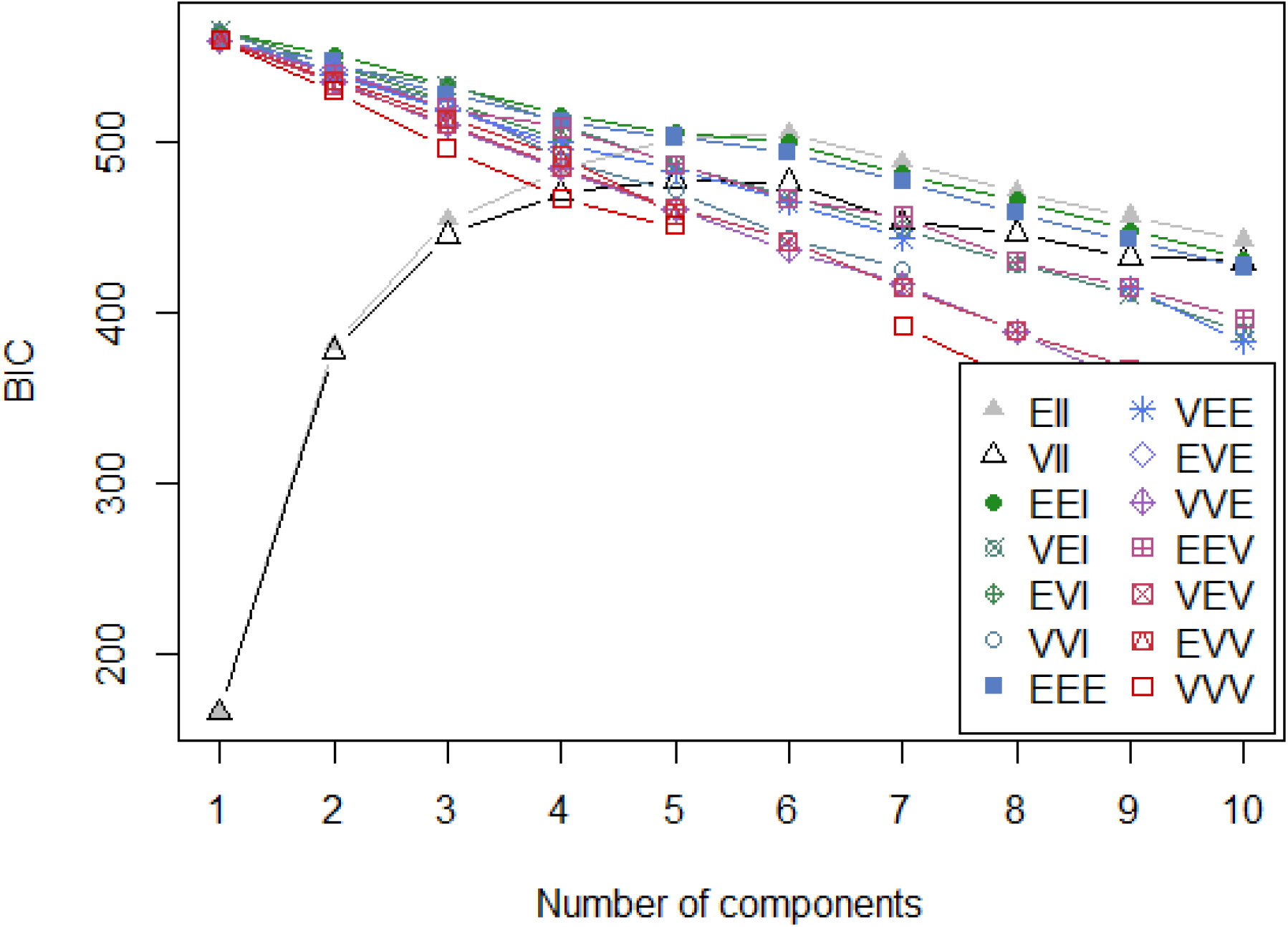
BIC scores of the *H. versicolor* NMM analysis for 1 to 10 components with individual lines representing the model classes fit by NMM specifying the shape, volume, and orientation of the mixture (see mclust package for more details).

**Supplemental Figure 10.**
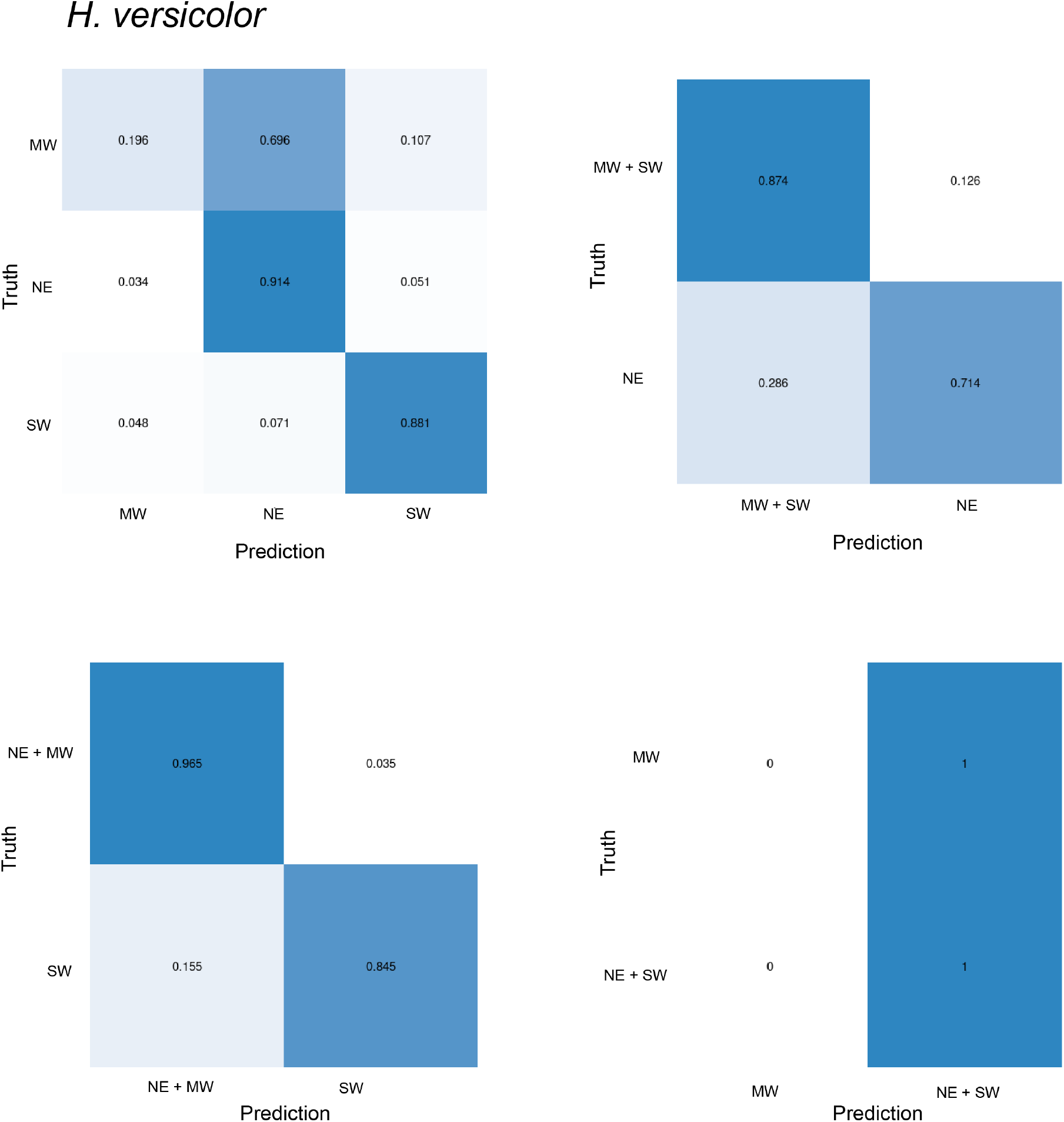
Confusion matrices of true and predicted components when components were pre-defined by *H. versicolor* mitochondrial lineage. The first confusion matrix demonstrates the precision for each component when all lineages are included, and the other matrices demonstrate the precision when two lineages are collapsed.

**Supplemental Figure 11.**
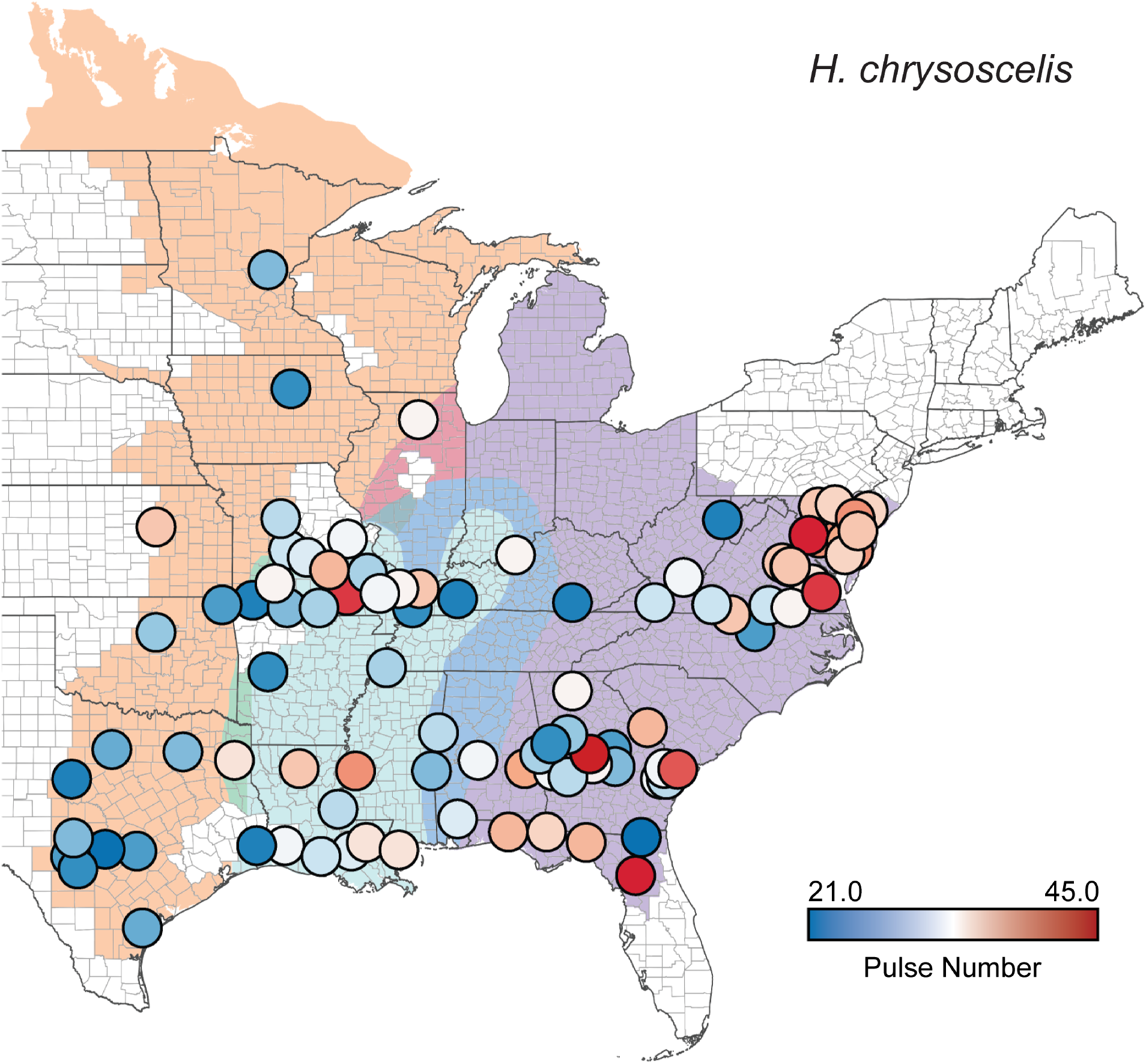
Call pulse number (total number of pulses in a call) in *H. chrysoscelis*. Circles represent averages for a single population. Points are colored from slow pulse number (blue) to high pulse number (red). Background colors correspond to mitochondrial lineages from Fig. 1.

**Supplemental Figure 12.**
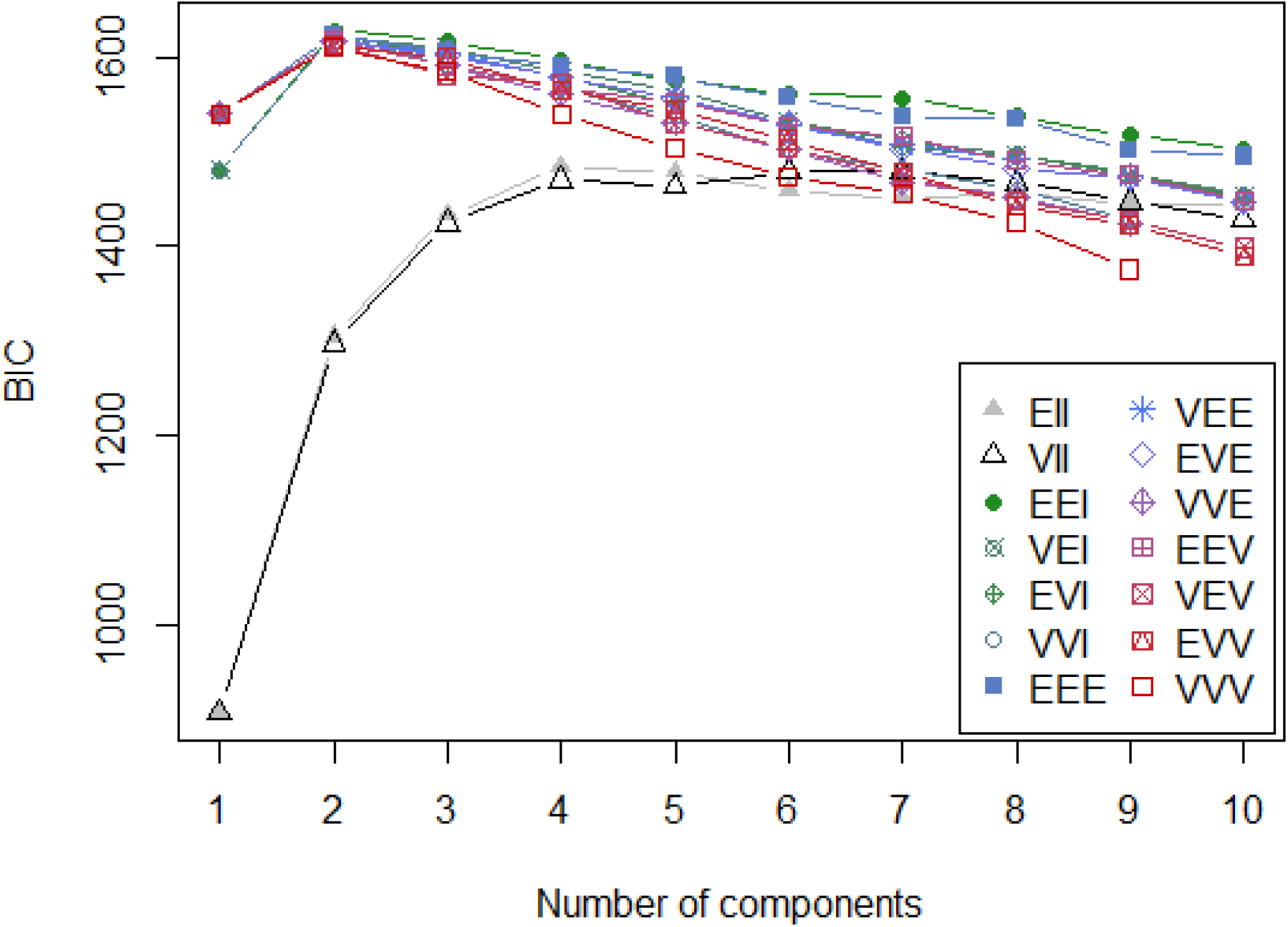
BIC scores of the *H. chrysoscelis* NMM analysis for 1 to 10 components with individual lines representing the model classes fit by NMM specifying the shape, volume, and orientation of the mixture (see mclust package for more details).

**Supplemental Figure 13.**
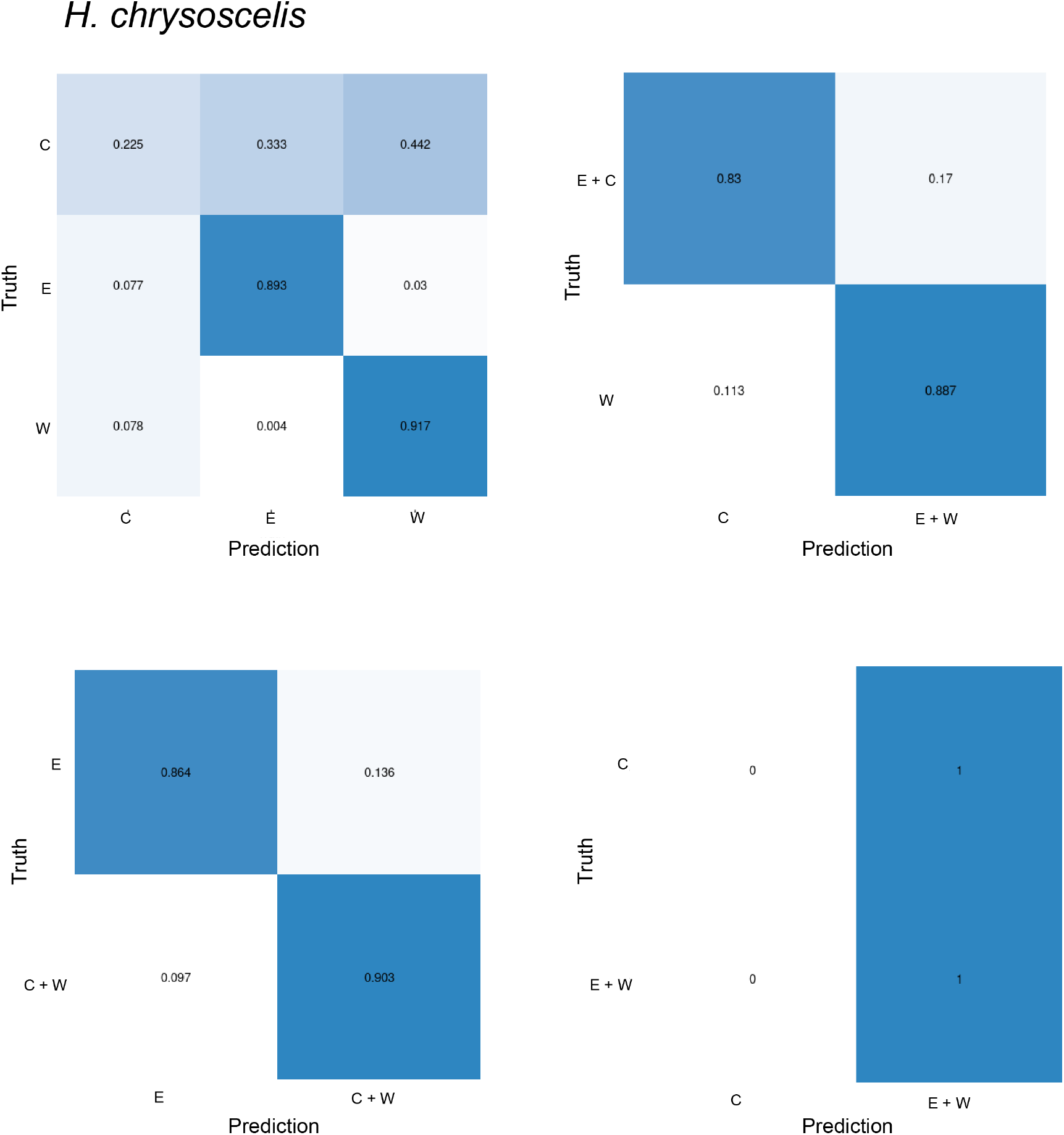
Confusion matrices of true and predicted components when components were pre-defined by *H. chrysoscelis* Selective regimes determined by PhylogeneticEM (Fig. 4a). The first confusion matrix demonstrates the precision for each component when all regimes are included, and the other matrices demonstrate the precision when two regimes are collapsed.

## References

1. Adams, K. L. and J. F. Wendel. 2005. Polyploidy and genome evolution in plants. Current Opinion in Plant Biology 8:135–141.

2. Angert, A. L., M. G. Bontrager, and J. Ågren. 2020. What do we really know about adaptation at range edges? Annual Review of Ecology, Evolution, and Systematics 51:341–361.

3. Austin, J. D., S. C. Lougheed, and P. T. Boag. 2004. Discordant temporal and geographic patterns in maternal lineages of eastern north american frogs, *Rana catesbeiana* (*Ranidae*) and *Pseudacris crucifer* (*Hylidae*). Molecular Phylogenetics and Evolution 32:799–816.

4. Austin, J. D., S. C. Lougheed, L. Neidrauer, A. A. Chek, and P. T. Boag. 2002. Cryptic lineages in a small frog: the post-glacial history of the spring peeper, *Pseudacris crucifer* (*Anura*: *Hylidae*). Molecular Phylogenetics and Evolution 25:316–329.

5. Avise, J. C. 2000. Phylogeography: the history and formation of species. Harvard university press.

6. Bagley, J. C., M. Sandel, J. Travis, M. d. L. Lozano-Vilano, and J. B. Johnson. 2013. Paleoclimatic modeling and phylogeography of least killifish, *Heterandria formosa*: insights into pleistocene expansion-contraction dynamics and evolutionary history of north american coastal plain freshwater biota. BMC Evolutionary Biology 13:223.

7. Baker, H. G. 1967. Support for baker’s law-as a rule. Evolution 21:853–856.

8. Baldwin, S. J. and B. C. Husband. 2011. Genome duplication and the evolution of conspecific pollen precedence. Proceedings. Biological Sciences / the Royal Society 278:2011–2017.

9. Barker, M. S., B. C. Husband, and J. C. Pires. 2016. Spreading winge and flying high: The evolutionary importance of polyploidy after a century of study. American Journal of Botany 103:1139–1145.

10. Barrow, L. N., A. T. Bigelow, C. A. Phillips, and E. M. Lemmon. 2015. Phylogeographic inference using bayesian model comparison across a fragmented chorus frog species complex. Molecular Ecology 24:4739–4758.

11. Bartlein, P., K. Anderson, P. Anderson, M. Edwards, C. Mock, R. Thompson, R. Webb, T. Webb III, and C. Whitlock. 1998. Paleoclimate simulations for north america over the past 21,000 years. Quaternary science reviews 17:549–585.

12. Bastide, P., C. Ańe, S. Robin, and M. Mariadassou. 2018a. Inference of adaptive shifts for multivariate correlated traits. Systematic Biology 67:662–680.

13. Bastide, P., C. Soĺıs-Lemus, R. Kriebel, K. William Sparks, and C. Ańe. 2018b. Phylogenetic comparative methods on phylogenetic networks with reticulations. Systematic Biology 67:800–820.

14. Blair, W. 1965. Amphibian speciation. Pages 543–556 in The Quaternary of the U.S. (H. E. Wright Jr. and D. G. Frey, eds.). Princeton University Press, Princeton, New Jersey, USA.

15. Bogart, J. P. and K. Bi. 2013. Genetic and genomic interactions of animals with different ploidy levels. Cytogenetic and Genome Research 140:117–136.

16. Bogart, J. P., P. Burgess, and J. Fu. 2020. Revisiting the evolution of the north american tetraploid treefrog (*Hyla versicolor*). Genome 63:547–560.

17. Booker, W. W., H. C. Gerhardt, A. R. Lemmon, M. B. Ptacek, A. T. Hassinger, J. Schul, and E. M. Lemmon. 2022. The complex history of genome duplication and hybridization in north american gray treefrogs. Molecular biology and evolution 39:msab316.

18. Bouckaert, R., J. Heled, D. Kühnert, T. Vaughan, C.-H. Wu, D. Xie, M. A. Suchard, A. Rambaut, and A. J. Drummond. 2014. BEAST 2: a software platform for bayesian evolutionary analysis. PLoS Computational Biology 10:e1003537.

19. Bretagnolle, F. and R. Lumaret. 1995. Bilateral polyploidization inDactylis glomerata l. subsp.lusitanica: occurrence, morphological and genetic characteristics of first polyploids. Euphytica 84:197–207.

20. Brochmann, C., A. K. Brysting, I. G. Alsos, L. Borgen, H. H. Grundt, A.-C. Scheen, and R. Elven. 2004. Polyploidy in arctic plants. Biological Journal of the Linnean Society 82:521–536.

21. Burbrink, F. T. 2002. Phylogeographic analysis of the cornsnake (*Elaphe guttata*) complex as inferred from maximum likelihood and bayesian analyses. Molecular Phylogenetics and Evolution 25:465–476.

22. Burton, R. S., R. J. Pereira, and F. S. Barreto. 2013. Cytonuclear genomic interactions and hybrid breakdown. Annual Review of Ecology, Evolution, and Systematics 44:281–302.

23. Cadena, C. D., F. Zapata, and I. Jiménez. 2018. Issues and perspectives in species delimitation using phenotypic data: Atlantean evolution in darwin’s finches. Systematic Biology 67:181–194.

24. Comai, L. 2005. The advantages and disadvantages of being polyploid. Nature Reviews. Genetics 6:836–846.

25. Curry, B. B., D. A. Grimley, and E. D. McKay. 2011. Quaternary glaciations in illinois. Pages 467–487 *in* Quaternary Glaciations Extent and Chronology A Closer Look vol. 15 of Developments in quaternary sciences. Elsevier.

26. David, K. T. 2022. Global gradients in the distribution of animal polyploids. Proceedings of the National Academy of Sciences 119:e2214070119.

27. De Smet, R., K. L. Adams, K. Vandepoele, M. C. E. Van Montagu, S. Maere, and Y. Van de Peer. 2013. Convergent gene loss following gene and genome duplications creates single-copy families in flowering plants. Proceedings of the National Academy of Sciences of the United States of America 110:2898–2903.

28. Delcourt, H. and P. Delcourt. 1984. Ice age haven for hardwoods. Natural History 93:22–25.

29. Drummond, A. J. and A. Rambaut. 2007. BEAST: Bayesian evolutionary analysis by sampling trees. BMC Evolutionary Biology 7:214.

30. Edwards, S. V. 2009. Is a new and general theory of molecular systematics emerging? Evolution 63:1–19.

31. Fontanella, F. M., C. R. Feldman, M. E. Siddall, and F. T. Burbrink. 2008. Phylogeography of *Diadophis punctatus*: extensive lineage diversity and repeated patterns of historical demography in a trans-continental snake. Molecular Phylogenetics and Evolution 46:1049–1070.

32. Freeling, M. and B. C. Thomas. 2006. Gene-balanced duplications, like tetraploidy, provide predictable drive to increase morphological complexity. Genome research 16:805–814.

33. Fullerton, D. S. 1986. Stratigraphy and correlation of glacial deposits from indiana to new york and new jersey. Quaternary science reviews 5:23–37.

34. Fusco, D., L. Grassi, B. Bassetti, M. Caselle, and M. Cosentino Lagomarsino. 2010. Ordered structure of the transcription network inherited from the yeast whole-genome duplication. BMC Systems Biology 4:1–9.

35. Gerhardt, H. 1994. Reproductive character displacement of female mate choice in the grey treefrog, *Hyla* chrysoscelis. Animal Behaviour 47:959–969.

36. Gerhardt, H. 1999. Reproductive character displacement and other sources of selection on acoustic communication systems. The design of animal communication. MIT Press, Cambridge, MA Pages 515–534.

37. Gerhardt, H. C. 1974. Mating call differences between eastern and western populations of the treefrog *Hyla* chrysoscelis. Copeia 1974:534.

38. Gerhardt, H. C. 1978. Temperature coupling in the vocal communication system of the gray tree frog, *Hyla versicolor*. Science 199:992–994.

39. Gerhardt, H. C. 1991. Female mate choice in treefrogs: static and dynamic acoustic criteria. Animal Behaviour 42:615–635.

40. Gerhardt, H. C. 2005. Advertisement-call preferences in diploid-tetraploid treefrogs (*Hyla chrysoscelis* and *Hyla versicolor*): implications for mate choice and the evolution of communication systems. Evolution 59:395–408.

41. Gerhardt, H. C. 2013. Geographic variation in acoustic communication: Reproductive character displacement and speciation. Evolutionary Ecology Research 15:605–632.

42. Gerhardt, H. C., M. L. Dyson, and S. D. Tanner. 1996. Dynamic properties of the advertisement calls of gray tree frogs: patterns of variability and female choice. Behavioral Ecology 7:7–18.

43. Gerhardt, H. C. and F. Huber. 2002. Acoustic communication in insects and anurans: common problems and diverse solutions. University of Chicago Press.

44. Gerhardt, H. C., M. B. Ptacek, L. Barnett, and K. G. Torke. 1994. Hybridization in the diploid-tetraploid treefrogs *Hyla chrysoscelis* and *Hyla versicolor*. Copeia 1994:51.

45. Gerhardt, H. C., M. A. Tucker, A. von Twickel, and W. Walkowiak. 2022. Anuran vocal communication: effects of genome size, cell number and cell size. Brain, Behavior and Evolution 96:137–146.

46. Gout, J.-F. and M. Lynch. 2015. Maintenance and loss of duplicated genes by dosage subfunctionalization. Molecular Biology and Evolution 32:2141–2148.

47. Gregory, T. R. and B. K. Mable. 2005. Polyploidy in animals. Pages 427–517 *in* The evolution of the genome (T. R. Gregory, ed.). Elsevier.

48. Griffin, A. R., T. D. Vuong, R. E. Vaillancourt, J. L. Harbard, C. E. Harwood, C. Q. Nghiem, and H. H. Thinh. 2012. The breeding systems of diploid and neoautotetraploid clones of *Acacia mangium* willd. in a synthetic sympatric population in vietnam. Sexual plant reproduction 25:257–265.

49. Harper, R. M. 1914. Geography and vegetation of northern florida. Florida Bureau of Geological Survey Annual Report 6:163–437.

50. Hegarty, M. and S. Hiscock. 2007. Polyploidy: doubling up for evolutionary success. Current Biology 17:R927–R929.

51. Holloway, A. K., D. C. Cannatella, H. C. Gerhardt, and D. M. Hillis. 2006. Polyploids with different origins and ancestors form a single sexual polyploid species. The American Naturalist 167:E88–101.

52. Husband, B., B. Ozimec, S. Martin, and L. Pollock. 2008. Mating consequences of polyploid evolution in flowering plants: Current trends and insights from synthetic polyploids. International journal of plant sciences 169:195–206.

53. Husband, B. C., S. J. Baldwin, and H. A. Sabara. 2016. Direct vs. indirect effects of whole-genome duplication on prezygotic isolation in *Chamerion angustifolium*: Implications for rapid speciation. American Journal of Botany 103:1259–1271.

54. Keller, M. J. and H. C. Gerhardt. 2001. Polyploidy alters advertisement call structure in gray treefrogs. Proceedings. Biological Sciences / the Royal Society 268:341–345.

55. Kirkpatrick, M. and N. H. Barton. 1997. Evolution of a species’ range. The American Naturalist 150:1–23.

56. Lemmon, A. R., S. A. Emme, and E. M. Lemmon. 2012. Anchored hybrid enrichment for massively high-throughput phylogenomics. Systematic Biology 61:727–744.

57. Lemmon, A. R. and E. M. Lemmon. 2008. A likelihood framework for estimating phylogeographic history on a continuous landscape. Systematic Biology 57:544–561.

58. Lemmon, E. M. 2009. Diversification of conspecific signals in sympatry: geographic overlap drives multidimensional reproductive character displacement in frogs. Evolution 63:1155–1170.

59. Lemmon, E. M. and A. R. Lemmon. 2010. Reinforcement in chorus frogs: lifetime fitness estimates including intrinsic natural selection and sexual selection against hybrids. Evolution 64:1748–1761.

60. Lemmon, E. M., A. R. Lemmon, and D. C. Cannatella. 2007a. Geological and climatic forces driving speciation in the continentally distributed trilling chorus frogs (*Pseudacris*). Evolution 61:2086–2103.

61. Lemmon, E. M., A. R. Lemmon, J. T. Collins, J. A. Lee-Yaw, and D. C. Cannatella. 2007b. Phylogeny-based delimitation of species boundaries and contact zones in the trilling chorus frogs (*Pseudacris*). Molecular Phylogenetics and Evolution 44:1068–1082.

62. Leroy, A. and P. Rousseeuw. 1987. Robust regression and outlier detection. Wiley Series in Probability and Mathematical Statistics, New York: Wiley, 1987 .

63. Levin, D. A. 1975. Minority cytotype exclusion in local plant populations. Taxon 24:35.

64. Levin, D. A. 2003. The cytoplasmic factor in plant speciation. Systematic Botany 28:5–11.

65. Li, Z., M. T. McKibben, G. S. Finch, P. D. Blischak, B. L. Sutherland, and M. S. Barker. 2021. Patterns and processes of diploidization in land plants. Annual Review of Plant Biology 72:387–410.

66. Love, E. K. and M. A. Bee. 2010. An experimental test of noise-dependent voice amplitude regulation in cope’s grey treefrog (*Hyla* chrysoscelis). Animal behaviour 80:509–515.

67. Lynch, M. and J. S. Conery. 2000. The evolutionary fate and consequences of duplicate genes. Science 290:1151–1155.

68. Lynch, M. and A. Force. 2000. The probability of duplicate gene preservation by subfunctionalization. Genetics 154:459–473.

69. MacArthur, R. H. 1972. Geographical ecology: patterns in the distribution of species. Harper Row.

70. Maherali, H., A. E. Walden, and B. C. Husband. 2009. Genome duplication and the evolution of physiological responses to water stress. The New Phytologist 184:721–731.

71. Marhold, K. and J. Lihová. 2006. Polyploidy, hybridization and reticulate evolution: lessons from the *Brassicaceae*. Plant Systematics and Evolution 259:143–174.

72. Martin, S. L. and B. C. Husband. 2012. Whole genome duplication affects evolvability of flowering time in an autotetraploid plant. Plos One 7:e44784.

73. Maxson, L., E. Pepper, and R. D. Maxson. 1977. Immunological resolution of a diploid-tetraploid species complex of tree frogs. Science 197:1012–1013.

74. McClintock, B. 1984. The significance of responses of the genome to challenge. Science 226:792–801.

75. McLachlan, J. S., J. S. Clark, and P. S. Manos. 2005. Molecular indicators of tree migration capacity under rapid climate change. Ecology 86:2088–2098.

76. Means, D. 1977. Aspects of the significance to terrestrial vertebrates of the apalachicola river drainage basin, florida. Florida Marine Research Publications 26:37–57.

77. Means, D. B. and K. L. Krysko. 2001. Biogeography and pattern variation of kingsnakes, lampropeltis getula, in the apalachicola region of florida. CHOREGIA Pages 1–9.

78. Miller, J. and D. Venable. 2000. Polyploidy and the evolution of gender dimorphism in plants. Science 289:2335–2338.

79. Muir, C. D. and M. W. Hahn. 2015. The limited contribution of reciprocal gene loss to increased speciation rates following whole-genome duplication. The American Naturalist 185:70–86.

80. Neill, W. 1957. Historical biogeography of present day florida: Bulletin of the florida state museum. Bulletin of the Florida State Museum, Biological Sciences 2:175–220.

81. Novikova, P. Y., I. G. Brennan, W. Booker, M. Mahony, P. Doughty, A. R. Lemmon, E. Moriarty Lemmon, J. D. Roberts, L. Yant, Y. Van de Peer, J. S. Keogh, and S. C. Donnellan. 2020. Polyploidy breaks speciation barriers in australian burrowing frogs *Neobatrachus*. PLoS Genetics 16:e1008769.

82. Novikova, P. Y., N. Hohmann, and Y. Van de Peer. 2018. Polyploid *Arabidopsis* species originated around recent glaciation maxima. Current Opinion in Plant Biology 42:8–15.

83. Oswald, B. P. and S. L. Nuismer. 2011. Neopolyploidy and diversification in *Heuchera grossulariifolia*. Evolution 65:1667–1679.

84. Otto, S. P. 2007. The evolutionary consequences of polyploidy. Cell 131:452–462.

85. Otto, S. P. and J. Whitton. 2000. Polyploid incidence and evolution. Annual Review of Genetics 34:401–437.

86. Pannell, J. R. and S. C. Barrett. 1998. Baker’s law revisited: reproductive assurance in a metapopulation. Evolution 52:657–668.

87. Parisod, C., R. Holderegger, and C. Brochmann. 2010. Evolutionary consequences of autopolyploidy. The New Phytologist 186:5–17.

88. Parks, C. R., J. F. Wendel, M. M. Sewell, and Y.-L. Qiu. 1994. The significance of allozyme variation and introgression in the l iriodendron tulipifera complex (*Magnoliaceae*). American Journal of Botany 81:878–889.

89. Peel, D. and G. MacLahlan. 2000. Finite mixture models. John & Sons.

90. Porturas, L. D. and K. A. Segraves. 2020. Whole genome duplication does not promote common modes of reproductive isolation in *Trifolium pratense*. American Journal of Botany 107:833–841.

91. Ptacek, M. B., H. C. Gerhardt, and R. D. Sage. 1994. Speciation by polyploidy in treefrogs: Multiple origins of the tetraploid, *Hyla versicolor*. Evolution 48:898.

92. Ralin, D. 1978. ”resolution” of diploid-tetraploid tree frogs. Science 202:335–336.

93. Ralin, D. B. 1976. Comparative hybridization of a diploid-tetraploid cryptic species pair of treefrogs. Copeia 1976:191.

94. Ramsey, J. 2011. Polyploidy and ecological adaptation in wild yarrow. Proceedings of the National Academy of Sciences of the United States of America 108:7096–7101.

95. Rice, A., P. S^̌^marda, M. Novosolov, M. Drori, L. Glick, N. Sabath, S. Meiri, J. Belmaker, and I. Mayrose. 2019. The global biogeography of polyploid plants. Nature Ecology & Evolution 3:265–273.

96. Rissler, L. J. and W. H. Smith. 2010. Mapping amphibian contact zones and phylogeographical break hotspots across the united states. Molecular Ecology 19:5404–5416.

97. Scrucca, L., M. Fop, T. B. Murphy, and A. E. Raftery. 2016. mclust 5: clustering, classification and density estimation using gaussian finite mixture models. The R journal 8:289.

98. Sharbrough, J., J. L. Conover, J. A. Tate, J. F. Wendel, and D. B. Sloan. 2017. Cytonuclear responses to genome doubling. American Journal of Botany 104:1277–1280.

99. Sloan, D. B., J. C. Havird, and J. Sharbrough. 2017. The on-again, off-again relationship between mitochondrial genomes and species boundaries. Molecular Ecology 26:2212–2236.

100. Soltis, D. E., A. B. Morris, J. S. McLachlan, P. S. Manos, and P. S. Soltis. 2006. Comparative phylogeography of unglaciated eastern north america. Molecular Ecology 15:4261–4293.

101. Stamatakis, A. 2014. RAxML version 8: a tool for phylogenetic analysis and post-analysis of large phylogenies. Bioinformatics 30:1312–1313.

102. Stebbins, G. L. 1984. Polyploidy and the distribution of the arctic-alpine flora: new evidence and a new approach. Bot. helvetica 94:1–13.

103. Stebbins, G. L. 1985. Polyploidy, hybridization, and the invasion of new habitats. Annals of the Missouri Botanical Garden 72:824.

104. Swenson, N. G. and D. J. Howard. 2005. Clustering of contact zones, hybrid zones, and phylogeographic breaks in north america. The American Naturalist 166:581–591.

105. Toews, D. P. L. and A. Brelsford. 2012. The biogeography of mitochondrial and nuclear discordance in animals. Molecular Ecology 21:3907–3930.

106. Tucker, M. A. and H. C. Gerhardt. 2012. Parallel changes in mate-attracting calls and female preferences in autotriploid tree frogs. Proceedings. Biological Sciences / the Royal Society 279:1583–1587.

107. Ueda, H. 1993. Mating calls in autotriploid and autotetraploid males in *Hyla japonica*. Sci. Rep. Lab Amphibian Biol. Hiroshima Univ., 12:177–189.

108. Van de Peer, Y., E. Mizrachi, and K. Marchal. 2017. The evolutionary significance of polyploidy. Nature Reviews. Genetics 18:411–424.

109. Welch, A. M. 2003. Genetic benefits of a female mating preference in gray tree frogs are context-dependent. Evolution 57:883–893.

110. Welch, A. M., R. D. Semlitsch, and H. C. Gerhardt. 1998. Call duration as an indicator of genetic quality in male gray tree frogs. Science 280:1928–1930.

111. Wells, K. D. and T. L. Taigen. 1986. The effect of social interactions on calling energetics in the gray treefrog (*Hyla versicolor*). Behav Ecol Sociobiol 19:9–18.

112. Werth, C. R. and M. D. Windham. 1991. A model for divergent, allopatric speciation of polyploid *Pteridophytes* resulting from silencing of duplicate-gene expression. The American Naturalist 137:515–526.

113. Yao, Y., L. Carretero-Paulet, and Y. Van de Peer. 2019. Using digital organisms to study the evolutionary consequences of whole genome duplication and polyploidy. PLoS One 14:e0220257.

114. Zeitler, L., C. Parisod, and K. J. Gilbert. 2022. Purging due to self-fertilization does not prevent accumulation of expansion load. bioRxiv Pages 2022–12.

